# Bisphenol-A mediated ubiquitinome alteration triggers PPAR-alpha ubiquitination, affecting trophoblast cell migration

**DOI:** 10.64898/2026.05.07.723151

**Authors:** Ankit Biswas, Sandhini Saha, Debapriyo Sarmadhikari, Krishna Singh Bisht, Shailendra Asthana, Tushar Kanti Maiti

## Abstract

Pregnant women are frequently exposed to various endocrine-disrupting chemicals (EDCs), such as bisphenol A (BPA), causing harm to both the developing placenta and fetus. BPA can promote placental dysfunction by altering key cellular processes such as differentiation, invasion, and migration in trophoblast cells. These cellular processes are also tightly managed by the ubiquitin proteasomal system via maintenance of the ubiquitinated protein pool. However, the BPA-mediated dysregulation of this ubiquitin proteasomal homeostasis is poorly understood. Therefore, we identified 19 deubiquitinases (DUBs) and a dynamic ubiquitinome profile of extravillous trophoblast cells (HTR8/SVneo), which reduced trophoblast cell migration post-BPA exposure. Further investigation using an integrated substrate-ligase-deubiquitinase network shows that BPA binding to PPAR-alpha or indirect regulation of its E3 Ligase MuRF1 and DUB USP5 via BPA resulted in hyper-ubiquitination of PPAR-alpha, triggering its nuclear localization. In the nucleus, the ubiquitinated PPAR-alpha can deregulate its migration-associated target gene expression, causing a reduction in the migration of HTR8/SVneo cells. This physiological alteration of extravillous trophoblast cells (EVTs) through BPA can disrupt placental homeostasis. Hence, we assumed that BPA-induced cellular alteration in EVTs can promote placental defects, which might contribute to adverse pregnancy outcomes.

## Introduction

The placenta is an endocrine organ that acts as a conduit between mother and fetus, undertaking circulatory, endocrine, and immune functions during pregnancy [1] . It originates from the trophectoderm of the blastocyst, containing stem cells known as cytotrophoblast, which differentiate into extravillous trophoblasts (EVTs) and syncytiotrophoblasts (SCTs) by invasion and syncytialization [2] Syncytiotrophoblasts produce hormones like human chorionic gonadotropin (hCG), progesterone, estrogen, and placental lactogens [3], which influence physiological functions such as migration, invasion, and differentiation of extravillous trophoblast cells [4] for the maintenance of a suitable uterine environment required for a healthy pregnancy. Trophoblast cells maintain this physiological homeostasis by expressing abundant hormone receptors, which increases their vulnerability to Endocrine Disrupting Chemicals (EDCs) [5]. EDCs are potent, synthetic, exogenous agents that can alter normal hormonal signalling in the human body, affecting reproduction, homeostasis, and developmental processes [6]. EDCs like phthalates, parabens, and bisphenols are predominantly present in everyday personal care products and plastic goods. During pregnancy, a woman can be exposed to these EDCs more frequently and easily, causing damage to both the developing placenta and fetus [7].

Bisphenol-A (BPA) is one such endocrine-disrupting chemical, which is known to cross the placental barrier and induce damage on both the placenta and fetus [8, 9]. BPA acts as a xenoestrogen mimicking hormonal signal which triggers oxidative stress and alters signalling pathways affecting trophoblast fusion, migration, invasion, and apoptosis [10]. Higher prenatal exposure to BPA is associated with the risk of various pregnancy complications like preeclampsia [11, 12], preterm birth [13], fetal growth restriction [14], and recurrent miscarriage [15]. However, the underlying molecular mechanism of BPA toxicity leading to these pregnancy complications is still unclear. Abnormal placentation is linked with various pregnancy complications [16]. Thus, many studies have explored the impact of BPA on placenta formation and function by using either trophoblast-derived cell lines such as BeWo, JEG3, and HTR-8/SVneo cells or, via primary trophoblasts and animal models. Based on the exposure concentration, BPA can elicit different physiological effects in HTR8/SVneo cells by altering DNA methylation [17], migration [18] and MAPK-PI3K signalling pathways [19]. Similarly, BeWo cells undergo apoptosis due to BPA-induced oxidative stress [20]. While JEG-3 cells show an anti-proliferative and pro-apoptotic phenotype [21] along with altered oestradiol metabolism [22] post-BPA exposure. BPA has also been shown to dysregulate the ubiquitin-proteasomal system in hippocampal neurons [23]. The ubiquitin signalling system has been shown to be a key regulator in cellular function, influencing protein turnover, cell cycle regulation, stress responses, and apoptosis [24,25]. Here, we hypothesize that BPA alters the balance of ubiquitination and de-ubiquitination processes, thereby influencing cellular signalling pathways that regulate trophoblast differentiation, migration, and invasion in placental cells. Although several comprehensive studies have examined the effects of BPA on placental function in both in-vitro and in-vivo models, the critical link between BPA-induced dysregulation of ubiquitin signalling and trophoblast biology remains unexplored.

In the present study, we elucidated the BPA-driven ubiquitin signalling regulation in extravillous trophoblast HTR8/SVneo cells by studying global ubiquitination, activity-based deubiquitinating enzymes profiling, and subsequently elucidating the crosstalk between ubiquitin ligases, deubiquitinase, and ubiquitinated substrates. Mass spectrometry-based global ubiquitination analysis identified PPAR-alpha’s ubiquitination at the ligand-binding domain and facilitated its nuclear localization. The higher expression of ubiquitin ligase MuRF1 and lower expression of deubiquitinase USP5 also corroborated the hyper-ubiquitination of PPAR-alpha. The ubiquitinated PPAR-alpha in the nucleus, activating or suppressing the expression of migration-associated genes, which orchestrated the reduced migration phenotype in HTR8/SVneo cells. This evidence underpins the hypothesis that BPA can cause hyper ubiquitination of PPAR-alpha, resulting in reduced migration of EVTs, disrupting the healthy functioning of the placenta, ultimately contributing towards various adverse pregnancy outcomes.

## Materials and methods

### Reagents

All general chemicals and reagents of LC-MS grade were purchased from Sigma-Aldrich (St. Louis, MO).

### Cell culture and Bisphenol-A treatment

HTR8/SVneo, a first-trimester extravillous trophoblast cell line, was kept in RPMI-1640 medium (SIGMA) that contains 2g/L sodium bicarbonate. 10% fetal bovine serum albumin (FBS) was supplemented in complete media. The cells were cultured in a CO_2_ incubator at 37 °C, with 5% CO2 and 85% relative humidity. Experiments were done using cells between 3-12 passages after thawing. Bisphenol A (Sigma Aldrich #133027) was dissolved in DMSO (Sigma) and added to the cell medium at a suitable dose, followed by incubation for 24 hours and 48 hours, after 4 hours of starvation in incomplete RPMI supplemented with 1% FBS. DMSO with less than 0.1% concentration in incomplete culture media was used as a control vehicle.

### Cell viability assay

MTT assay was used to measure the cell viability. Inside the living cell, the active enzymes in mitochondria reduce the MTT tetrazolium salts, which were further detected calorimetrically as a measure of cell viability. Phosphate-buffered saline (PBS) was used for the preparation of Thiazolyl Blue Tetrazolium Bromide (MTT) (Sigma) at a final concentration of 5 mg/ml. In a 96-well plate 1x10^4^ HTR8/SVneo cells per well were seeded in triplicates that contain 200 μL RPMI with 10% FBS and maintained 24-36 hours before BPA treatment with specific concentration (0-500 μM) for 24 and 48 hours. After the treatment was complete, in each well of 96-well plates 10 μL stock MTT solution was added and incubated for 4 hours at 37 °C. Then the media was discarded. To dissolve the reduced MTT crystals 200 μL 100% DMSO was added to each well and incubated for 20 min at 37 °C. The colorimetric intensity was measured using a microplate reader (Spectramax) at a wavelength of 562 nm.

### Cellular migration assay

3X10^5^ HTR-8/SVneo cells in 6-well plates were plated containing 2 ml culture medium and incubated for 36-48 h. After the cells were confluent, they were first starved for 4 hours in 1% FBS-supplemented RPMI and then incubated with 10 μM of BPA for 24 and 48 h. After the treatment was complete, a wound was generated by scraping off the cells with a sterile pipette tip and photographed every 12 hours for 2-3days. DMSO (0.1% in medium) was used as a control for BPA treatments. The scratch was photographed by removing unattached cells in the existing medium, and the uncovered area was measured at each time point using ImageJ software.

### Annexin-FITC/Apoptosis assay

12-well plate was seeded with 1X10^5^ HTR8/SVneo cells per well containing 1ml Media and maintained for 36-48 hours, then starved for 4 hours in 1% FBS-supplemented media followed by incubation in 10 μM BPA or 0.1% DMSO (control vehicle) for 24 and 48 hours. After complete incubation, the cells from each well were trypsinized and collected in different 1.5 ml MCTs and washed with 500 μL chilled PBS 2 times. Then the pellets were dissolved in the 1X binding buffer provided in the Bio-Vision Kit, followed by the addition of 5 μL of Annexin-FITC and PI dye provided in the kit. After 10 minutes of incubation, the samples were analyzed by flow cytometry using a BD FACS. For each sample, a minimum of 10,000 cells was analyzed. Based on the forward- and side-scatter properties, the cell population was gated. The auto-fluorescence of control cells was used to designate Cut-off values (i.e., vertical and horizontal lines on the resulting scatter plots). The FlowJo software package was used for the data analysis and represented the percentage of cells in each phase (healthy, early apoptosis, late apoptosis, necrosis). Positive control and negative control were prepared by subsequently giving a heat shock at 95 °C for 15 minutes and maintaining cells in complete RPMI during treatment, respectively.

### Immuno-fluorescent analysis of cells

12-well plates were seeded with 1X10^5^ Cells per well on cover slips having 1 ml medium and maintained for 24 hours then treated with 10 μM BPA for 24 hours as done earlier. Next, the media was discarded, and the coverslips were fixed with 4% paraformaldehyde for 10 minutes at RT, followed by three times washing with 1X PBS. The cover slips were transferred to a humidified chamber and blocked with 1% BSA in PBST (PBS and 0.1% Triton-X 100) for 1 hour, followed by incubation in 60 μL primary antibody (**Table S1**) diluted in 1% BSA in PBST for 1.5 hours. After incubation, the coverslips were washed with 1X PBS for 3 times and then 60 μL secondary antibody (Alexa fluor-594 goat anti-mouse lgG and Alexa fluor-684 goat anti-rabbit lgG, Thermo Fisher Scientific, USA), diluted 500 times in 1% BSA in PBST, was added and incubated for 1 hour and again washed with 1X PBS for 3 times. Then 60 μL DAPI from 5 μg/ml stock was added to each coverslip and incubated for 5 minutes and washed with 1X PBS 3 times. Then the coverslips were washed once in water and kept for drying. When completely dried, the coverslips were mounted in a glass slide using mounting reagent and kept overnight in the dark at RT to dry. Then stored in a slide box at -20 °C. Images were obtained in a confocal microscope (Leica TCS SP8) and processed in ImageJ software to analyze the colocalization of ubiquitin and PPAR-alpha in the nucleus.

### Immuno-precipitation

100 mm plates seeded with 70% confluent cells were treated with 10 μM BPA as described previously. After 24 hours of incubation the whole cell lysate was prepared in IP lysis buffer (25 mM Tris-HCl pH 7.4, 150 mM NaCl, 1 mM EDTA, 1% NP-40 and 5% glycerol) supplemented with a 1X halt-protease and phosphatase inhibitor cocktail. The total protein concentration was measured using BCA reagent kit (Thermo #23227) and 500 μg of protein form each condition were incubated with 5 μg of primary antibody (**Table S1**) for 2 hours at 4 °C. The antigen-antibody complex was then captured using protein-G dynabeads (Thermo, Cat#10003D) by incubating the beads overnight with antibody mixed protein lysate at 4 °C. Next day, the beads were washed 3 times with IP lysis buffer followed by a single wash with Milli-Q water and the enriched proteins were eluted two times by heating the dynabeads at 95 °C in 2X lamellae dye. The entire elution volume was subjected to SDS-PAGE and subsequently analyzed by Immuno-blotting using the corresponding antibodies.

### Immuno-blot analysis

In a 6 well plate 5X10^5^ Cells were seeded in each well that contains 2ml media and kept for 36-48 hours. The treatment was done as previously and after completion of treatment 100 μL RIPA containing 1X PPI was added to each well and scrapped. Different treatment condition was collected in different 1.5 ml MCT, then the MCT were incubated in ice for 30 mins with intermitted vortexing. Then the lysate was subjected to probe sonication followed by centrifugation at 16000 rpm for 45 mins at 4 °C. then the supernatant was collected in a different 1.5ml MCT and the protein concentration was measured by BCA reagent kit (Thermo #23227). Electrophoretic separation of equal amounts of the protein samples (25 μg) was done on a 12% sodium dodecyl sulphate polyacrylamide gel (SDS-PAGE) followed by transferred to a 0.45μ polyvinylidene fluoride (PVDF) membrane (Merk) and blocking with 5% skimmed milk in 0.1% PBST/TBST at room temperature for 1 h, the membranes were incubated overnight with primary antibodies (**Table S1**) at 4 °C. After incubation and washing with 0.1% Tris-buffered Saline Tween (TBST)-20, the membrane was incubated with secondary anti-rabbit or anti-mouse antibody (1:10000 & 1:10000 respectively) at room temperature for 1 h. After, washing with TBST, the membrane was developed using Immobilon Forte Western HRP Substrate (Millipore, MA, USA). ImageJ software was used for the densitometric quantification of the blots. The relative intensity of each band was normalized to that of GAPDH or β-actin, serving as loading controls for the same blot.

### qRT-PCR analysis

The total RNA was extracted from 10 μM BPA treated cells along with control using TRIzol reagent (Invitrogen). Briefly, the cells were collected in 500 μL of TRIzol reagent, and following phase separation with chloroform, the aqueous phase was collected. RNA precipitation was done by adding isopropanol, followed by washing with 75% ethanol, and resuspension in nuclease-free water. The quantity and quality of the extracted RNA were assessed using a NanoDrop spectrophotometer (Thermo #ND-ONE-W). cDNA was synthesized with 1 μg of RNA using the iScript cDNA Synthesis Kit (Bio-Rad #1708891) in a 20 μL reaction containing iScript Reverse Transcriptase and 5x Reaction Mix. The reaction was performed at 25 °C for 5 minutes, 46 °C for 20 minutes, and 95 °C for 1 minute. Next, for qRT-PCR, 1 μL of diluted cDNA was used in a reaction mixture of 20 μL with 10 μL of Universal SYBR Green Supermix (Bio-Rad #1725121), 0.4 μM forward and reverse gene-specific primers (**Table S2**) and GAPDH as the housekeeping control diluted in RNAase-free water. The qRT-PCR cycling conditions were initial denaturation at 95 °C for 3 minutes, followed by 40 cycles of 95 °C for 10 seconds and 60 °C for 30 seconds in QuantStudio 6 (Applied Biosystems). Data were analyzed using the ΔΔCt method, with GAPDH as the reference gene.

### In-silico BPA binding

#### Structure Retrieval and System Preparation

Since no crystal structure of apo PPAR-alpha has been reported in the Protein Data Bank (PDB), the structural model of PPAR-alpha was obtained from the AlphaFold database [26]. To identify the potential BPA binding site in PPAR-alpha, we utilized the crystal structure of PPAR-gamma in complex with BPA (PDB ID: 9F7W) [27] For ubiquitin docking studies with PPAR-alpha, the crystal structure of ubiquitin (PDB ID: 2L0T) [28] was used and to understand the dynamic motion of PPAR-alpha structure we have taken all available crystal sutures (69 crystals) in RCSB database. All protein structures were processed using the Protein Preparation Wizard in Maestro (Schrödinger Release 2023-3) [29]. Hydrogen atoms and bond orders were assigned using PRIME, followed by hydrogen-bond optimization and restrained energy minimization employing the OPLS4 force field [30–32].

#### Molecular Docking

Protein-ligand docking studies were carried out using AutoDock4.28 to predict the binding mode of BPA within PPAR-alpha. The PPAR-gamma-BPA complex (PDB ID: 9F7W) was used as a structural reference to define the docking site [33, 34]. The PPAR-alpha model and BPA ligand were prepared in AutoDock Tools (ADT) by adding polar hydrogens, merging nonpolar hydrogens, assigning Gasteiger charges, and defining ligand torsions [35]. The docking grid was focused on the putative BPA binding pocket of PPAR-alpha and for calculation we used Lamarckian Genetic Algorithm (LGA). A total of 10 ligand poses were obtained to further quantify the binding pose and mode [34]. For the protein-protein docking of ubiquitin with PPAR-alpha, the HDOCK standalone software was employed, following default scoring and refinement protocols [36].

#### Residue label stability prediction and post processing analysis

Structural stability of PPAR-alpha, both in BPA-bound and unbound forms, was evaluated using normal mode analysis (NMA) implemented via the ProDy plugin in VMD [37–40]. All post-processing analyses were performed in VMD, and the resulting plots were generated using XMGRACE.

### Activity-based deubiquitinating enzyme profiling

#### Sample preparation

3X10^6^ cells were seeded in 100 mm plates in replicates and maintained for 36-48 hours to reach the confluency at 70%. Then, treated with 10 μM BPA or 0.1% DMSO as mock control for 24 and 48 hours and finally harvested in DUBs lysis buffer (50mM Tris, 5mM MgCl2, 250mM Sucrose, 0.5% CHAPS, 0.1% NP-40, pH 7.4) containing 1mM PMSF. Incubated in ice for 30 mins and then probe sonication was done. After that, the lysate was centrifuged at 14,000 rpm for 45 mins at 4 °C and the supernatant was collected in a different MCT. The protein concentration of each lysate was quantified by a BCA reagent kit (Thermo) using BSA as a standard. Then, 500 μg and 1 mg of protein were aliquoted respectively for western blot and mass spectrometry experiments and respectively 40 μL and 80 μL of 25 μM HA-UbVMe probe in 50 mM sodium acetate, pH 4.5 was added to the lysate followed by addition of double volumes 50 mM sodium hydroxide (NaOH) compared to probe volumes and checked for pH 8 using pH paper. Then the mixture volumes were made up to 500 μL and 1ml respectively using DUBs lysis buffer so that the protein concentration would be 1mg/ml and incubated for 4 hours at RT. The anti-HA magnetic beads (Pierce, USA) were prepared by washing them with an increased volume of 0.05% TBST and then added to the probe incubated lysate premixed with 1X protein phosphatase inhibitor. The mixture was incubated overnight at 4 °C in a head-to-head rotator. The next day, the beads were washed first with 0.05% TBST followed by 800 μL ultrapure water for 2 times. Then the bound proteins were eluted in 30 μL of 5X lamellae dye by heating the beads at 95 °C for 5 minutes. For western blots, the lower mentioned protocol was used afterward. In-Gel digestion was done for mass spectrometry analysis

#### In-gel digestion and data acquisition

The eluted proteins were run on 12% SDS-PAGE gel at 100V to resolve only 40% of gel then properly stained and destained and further washed in autoclaved Milli-Q water to remove any residual acetic acid. Then the whole lane of each well was cut and distributed in different MCTs containing 1mm^3^ pieces of gel which were washed in 50% acetonitrile in 50 mM ammonium bicarbonate properly to remove any staining dye. Then, dehydrated in 100% acetonitrile followed by rehydration with 150 μL reduction solution (10mM DTT in 100mM ammonium bicarbonate) and incubated at 56 °C for 30 mins. Next 100 μL alkylation solution (20mM Iodoacetamide in 100mM ammonium bicarbonate) was added after removing the previous solution and incubated at RT for 30 mins. Then, 110 μL of 10 g/ml pierce trypsin (Thermo) was added to each tube and incubated at 37 °C for 18 hours. After the completion of digestion, the peptides were eluted in 50 μL extraction solution (60% acetonitrile and 0.1% formic acid) and peptides from a single lane were pooled in a single MCT and dried. Then, the peptides were zip-tipped using C-18 tips (Thermo) and again dried and stored at -80 °C. The digested peptides were vacuum dried and reconstituted in 40 μL of solvent A (2% (v/v) ACN, 0.1% (v/v) FA in water) and subjected to LC-MS/MS experiments using Sciex 5600+ Triple-TOF mass spectrometer coupled with ChromXP reversed-phase 3 m C18-CL trap column (350 μm X 0.5 mm, 120 Å, Eksigent, AB Sciex) and nanoViper C18 separation column (75 μm X 250 mm, 3 μm, 100 Å; Acclaim Pep Map, Thermo Scientific, USA) in Eksigent nanoLC (Ultra 2D plus) system. 5 μL from each sample were injected with 250 nL/min flow rate with an increasing linear gradient of solvent B (98% (v/v) ACN, 0.1% (v/v) FA) until 16 min for a total run time of 35 mins. The data acquisition was done with data-dependent acquisition (DDA) mode. Parent spectra were acquired with the scan range of 400-1250 m/z for top 25 ions with minimum intensity of 120 cps. The mass tolerance and charge state were set to 50 mDa and 2-5 respectively. Data-dependent acquisition experiments were set to obtain a high-resolution TOF-MS/MS scan Using Collision induced dissociation with pulser frequency of 14.980 kHz over a mass range of 100-1600 m/z. The number of cycle was 2583 with the accumulation time of 250 ms for TOF-MS and 100 msec for MS/MS. The mass spectrometry proteomics data have been deposited to the ProteomeXchange Consortium via the PRIDE partner repository with the dataset identifier PXD065005.

#### Data analysis

The generated chromatogram was searched in protein pilot (version 4.5, SCIEX) software using a mascot (version 2.3.02) search engine against SwissProt 57.15 database (20266 sequences after Homo sapiens taxonomy filter) using the following criteria: enzyme trypsin, Maximum missed cleavage 1, MS/MS fragment tolerance 0.2 Da, precursor tolerance 100 ppm and peptide charge 1+, 2+, and 3+ with Monoisotopic mass, No fixed modifications, and variable modifications of carbamidomethylation in cysteine (+57.02146 Da), deamination of NQ (+0.98416), UB-G-VMe (172.084792), UB-LRG-VMe (441.269967), UB-LRG-VMe(H) (442.277792), UB-VMe (173.0922617), UB-VMe-S(NH3+) (206.072513), UB-VMe-SH (205.0646688). The generated search file (.mrf) was exported in .csv format and deubiquitinase enzymes from the list of identified proteins were extracted for further visualization using UpSet plot and heatmap using UpSetR (v1.4.0) and pheatmap (v1.0.12) Package respectively in R (v4.3.1)

### Global ubiquitinome profiling

#### Sample preparation and tryptic digestion

Global ubiquitinome analysis was carried out using anti-diGly affinity enrichment coupled with LC-MS/MS. For this, HTR8/SVneo cells were incubated with 10 μM BPA for 24 hours, where cells were treated with a control vehicle and used as a control. After completion of 24 hour, cells were harvested and proteins were extracted by RIPA lysis buffer (Sigma, #R0278) supplemented with 1X halt-protease and phosphatase inhibitor cocktail (Thermo #1861282) with intermediate vortexing, followed by mild sonication. The lysate was centrifuged at 14,000 g for 30 min at 4 °C and the supernatant was collected. Six volumes of acetone were added to the lysates for protein precipitation. The protein precipitates were re-solubilized in 8 M urea and concentrations were estimated by the BCA method (Pierce BCA Protein Assay Kit, Thermo Scientific). From each condition, an equal amount of protein (6.5 mg) was incubated with 10 mM DTT (56 °C, 1 hour) for reduction and alkylated with 20 mM IAA (room temperature, 1 hour, dark). MS grade trypsin (Sigma Aldrich, USA) was added for digestion (trypsin: protein ratio of 1:10 (w/w) at pH 8, 37 °C for 36 hours) and reactions were quenched by adding 1% formic acid to the final volume after complete digestion. Following digestion, the resulting peptide mixtures were centrifuged at 14000g for 30 min to remove the undigested particles. The supernatant was collected and loaded onto the reversed-phase C18 Sep-Pak cartridges (Waters #WAT020515) for desalting and clean-up of peptides. The peptides were then vacuum dried and kept at -80 °C until enrichment.

#### k-εGG (diGly) peptide enrichment

diGly modified peptides enrichment was done by the di-glycyl-lysine Antibody (Lucerna technologies). Briefly, dried di-glycyl-lysine Antibody was reconstituted in 1 ml 0.2 M HEPES (pH 8.5) buffer and dried peptides were resuspended in IP lysis buffer (25 mM Tris-HCl pH 7.4, 150 mM NaCl, 1 mM EDTA, 1% NP-40 and 5% glycerol, Thermo Scientific). Each peptide (6.5 mg) was incubated with 30 μL of anti-diGly antibody overnight at 4 °C at an end-to-end rotor. Pre-activated and washed protein G beads were added to the peptide-antibody complex mixture after overnight incubation. Next, these beads were incubated with antibody bound peptide mixture at 4 °C overnight while rotating. After complete incubation, each falcon was centrifuged at 4000g for 5 min and the supernatant was removed. Beads were washed twice with 1 ml of PBS and once with 1 ml washing buffer (5% acetonitrile in PBS), and finally once in ultrapure water. Ubiquitinylated peptides were eluted by adding 0.3 ml of a solution containing 0.1% formic acid and 80% acetonitrile in water by boiling at 95 °C for 5min. A total of 10 elutions were collected in sample MCT for each sample and dried in a SpeedVac. Further the dried samples were desalted with C18 tips (Pierce, Thermo Fisher Scientific, USA) and reconstituted with solvent A (2% (v/v) acetonitrile, 0.1% (v/v) formic acid in water) for LC-MS/MS analysis.

#### Identification of diGly modified peptides

LC-MS/MS experiments were performed using Sciex 5600+ Triple-TOF mass spectrometer coupled with ChromXP reversed-phase 3 m C18-CL trap column (350 μm X 0.5 mm, 120 Å, Eksigent, AB Sciex) and nanoViper C18 separation column (75 μm X 250mm, 3 μm, 100 Å; Acclaim Pep Map, Thermo Scientific, USA) in Eksigent nanoLC (Ultra 2D plus) system. The binary mobile solvent system was used as follows: solvent A (2% (v/v) ACN, 0.1% (v/v) FA in water) and solvent B (98% (v/v) ACN, 0.1% (v/v) FA). The peptides were separated using a 60 min gradient with a total run time of 90 min at a flow rate of 300 nL/min. The MS data of each condition was acquired in IDA (information-dependent acquisition) with high sensitivity mode. Each cycle consisted of ∼250 and 100 ms acquisition time for MS1 (m/z 350-1250 Da) and MS/MS (100-1600 m/z) scans respectively, with a total cycle time of ∼2.8s. Each condition was run in triplicate. The mass spectrometry proteomics data have been deposited to the ProteomeXchange Consortium via the PRIDE partner repository with the dataset identifier PXD065000.

#### Ubiquitinome data analysis

The wiff spectral files were used to search the data in “ProteinPilot” software (version 4.5, SCIEX) using the Mascot algorithm (version 2.3.02) for the identification of the proteins against the SwissProt_57.15 database (20266 sequences after Homo sapiens taxonomy filter). To identify the ubiquitinated peptides the following search parameters were used: a) The proteolytic enzyme was set as trypsin with two maximum allowed missed cleavages; b) Peptide mass tolerance of 20 ppm and fragment mass tolerance of 0.2 Da; c) Carbamidomethylation of cysteine (+57.021464 Da), oxidation of methionine (+15.994915 Da), deamination of NQ (+0.984016) and Gly-Gly of Lysine (114.042927) were used as variable modifications. The searched list of proteins and peptides were then further sorted to extract Ubiquitin modified protein and peptides from non-modified fraction. The PEP and Prot score distribution were checked to measure the quality of the data followed by analysis of uniquely modified ubiquitinated proteome in BPA treated condition using Metascape (v3.5.20250701) webtool and R (v4.3.1) language.

#### Bioinformatic and statistical analysis

We build an integrated network of deubiquitinase, E3 ligase and the ubiquitinated substrate using information from Ubibrowser 2.0 database. First, we selected the putative targets of the identified deubiquitinases from our data and then overlapped with the uniquely ubiquitinated substrate identified in BPA treated condition. The putative E3 ligases of the common substrates were identified with a cut-off value of ≥ 0.8. The compiled network was then visualized using Cytoscape software (Version 3.8.2). The genes regulated by transcription factor PPAR-alpha, beta and gamma were identified using webtool TFLink (https://tflink.net/) based on only small-scale evidence. The pathway enrichment of common substrate proteins and unique genes regulated by PPAR-alpha was done using ShinyGO v0.77 (https://bioinformatics.sdstate.edu/go77/) platform. STRING (v12.0) database were employed to visualize the PPI network between genes uniquely regulated by PPAR-alpha.

Statistical analysis for all the experiments was performed using Prism 8 software (GraphPad Software, USA). For data plotting and visualization either Prism 8 or ggplot (3.5.1) were used. All data were obtained for at least three biological replicates and were expressed as mean ± SEM if not mentioned either. The significance of differences between control and BPA treatment groups were determined using student’s t-test or two-way ANOVA and P values *<* 0.05 were considered statistically significant and indicated by asterisks as follows: *P *<* 0.05, **P *<* 0.01, ***P *<* 0.001, ****P *<* 0.0001. All the graphical illustrations were made using BioRender (https://www.biorender.com/).

## Results

### BPA exposure reduces the trophoblast cell migration and alters global protein homeostasis

HTR8/SVneo cells are susceptible to external BPA exposure, which disrupts their various physiological functions like migration, invasion and apoptosis [17–19]. Thus, to identify the susceptible BPA dose and its subsequent effect in trophoblast cells, we followed the schematic as depicted in **Figure 1A**. MTT assay suggested no significant reduction in cell viability till 125 μM and 31.25 μM concentration after 24 and 48 hours of incubation, respectively (**Figure1B**). Thus, based on MTT assay and published literature [17–19], we selected 10 μM concentration for future experiments. The dose of 10 μM BPA showed a significant reduction in cell migration when exposed for 24 hours, followed by the scratch assay (**Figure 1C and Figure S1A**). But no significant changes were observed for 48 hours of exposure (**Figure 1C and Figure S1B**). We found that the expression of MMP2 and MMP9 was down-regulated significantly after 10 μM BPA exposure for 24 hours. However, consistent with the previous result, 48 hours of incubation showed no significant changes (**Figure 1D, E**). Apoptosis assay also showed no changes in percentage of healthy, early apoptotic, late apoptotic or necrotic cells when compared with control in case of both 24 hours (**Figure 1F and Figure S1C**) and 48 hours (**Figure 1G and Figure S1D**) of BPA administration.

**Figure 1:**
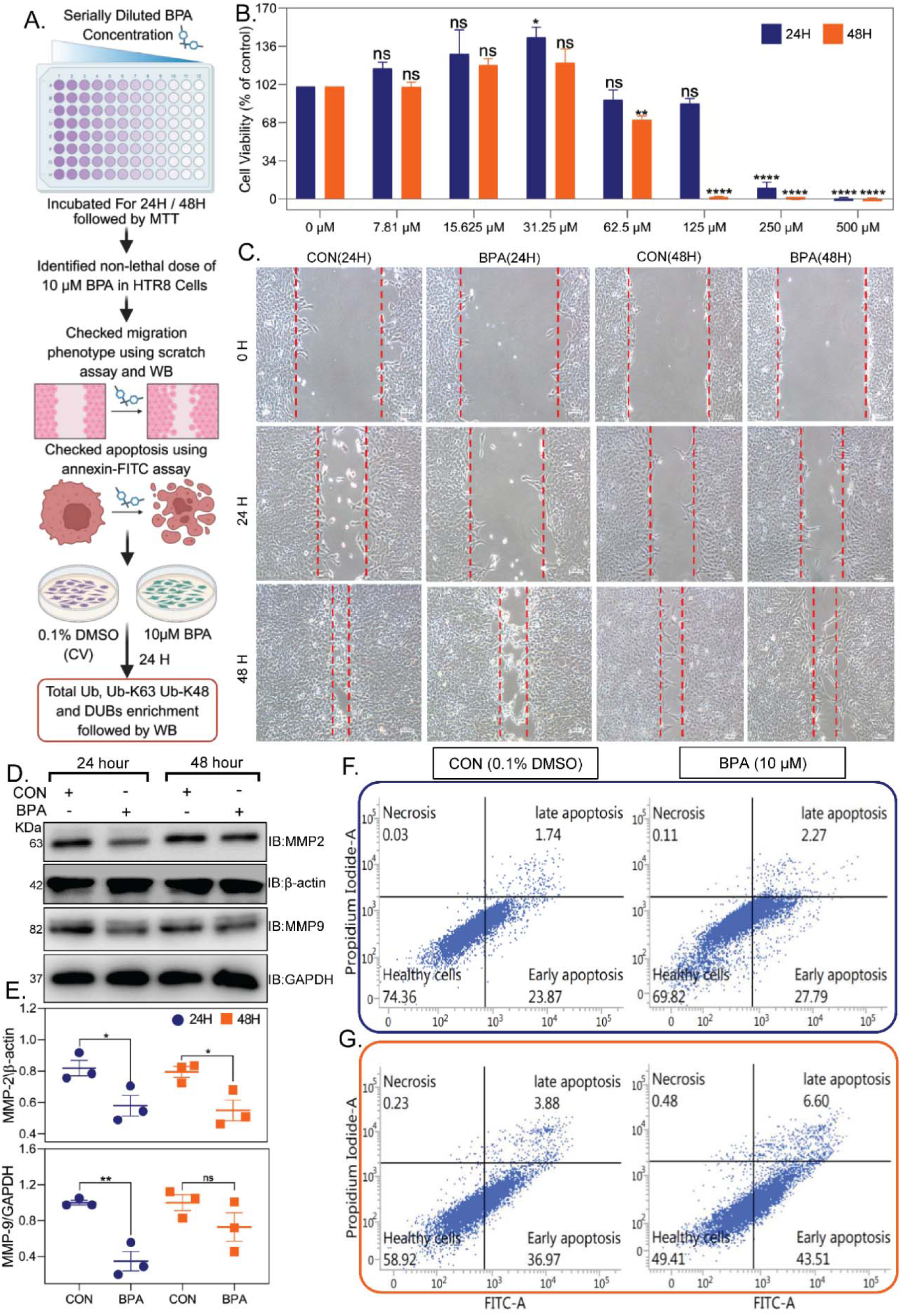
BPA exposure reduces cell migration in HTR8/SVneo cells. **A)** Schematic workflow describing the stepwise experiments done to determine the susceptible BPA dose and its subsequent effect on HTR8/SVneo cells. **B)** Cell viability was calculated as (abs of test*/*abs of control) ×100% for serially diluted BPA concentrations exposed for 24 hours (dark blue) and 48 hours (orange). Data are shown as mean ± SEM (N=3) and analyzed using ordinary one-way ANOVA, considering a p-value *<* 0.05 as significant. **C)**Phase-contrast images of HTR8/SVneo cells at 0, 24, and 48 hours after scratch injury, treated with 10 μM BPA or vehicle control for 24 or 48 hours. Wound closure was observed after 48 hours. Scale bar = 100 μm. **D)** Representative immunoblot for MMP2 and MMP9 expression after 24 and 48 hours of incubation with 10 μM BPA or control vehicle. **E)** Densitometric quantification of the blots, normalized using β-actin or GAPDH for MMP2 and MMP9, respectively, is represented as mean ± SEM of three independent experiments. Statistical significance was determined using Student’s t-test, with *p <* 0.05 considered significant (*P *<* 0.05, **P *<* 0.01). Representative plots for flow cytometry showing the apoptosis assay performed using Annexin V-FITC and propidium iodide (PI) staining after **F)** 24 hours and **G)** 48 hours of Control vehicle or 10 μM BPA treatment. Cells were initially gated on live, single cells (FSC-H vs. SSC-H), and then Annexin V and PI were used to distinguish early apoptotic (Annexin V+ PI-), late apoptotic (Annexin V+ PI+) and necrotic (Annexin V-PI+) populations. The percentage of cells in each quadrant was shown.

The findings indicate that increased BPA concentration and duration of exposure significantly reduce the cell viability, as described in previous literature [17]. Hence, long-term and chronic exposure to BPA could be detrimental to the placenta during pregnancy. As a xenoestrogen, BPA is known to impair migration and invasion [17, 18] of extravillous trophoblasts (EVTs). We found that 10 μM BPA exposed for 24 hours reduced the migration of HTR8/SVneo cells significantly but did not induce apoptosis. The findings suggest that low-dose BPA does not exert a direct cytotoxic effect but modulates its physiological functions. Migration is a crucial event in EVTs, initiating the structural remodelling of the uterine spiral artery via invading maternal decidua to secure placental blood flow essential for fetal development [2]. Migratory cells require a dynamic proteome, maintaining a fine balance between protein synthesis and degradation. This balance is tightly regulated by the ubiquitin-signalling system [24]. Thus, reduction of migration should also impair the ubiquitin-signalling network by altering protein ubiquitination and de-ubiquitination events. Therefore, it is pertinent to investigate the ubiquitination and de-ubiquitination status of HTR8/SVneo cells after BPA exposure.

Interestingly, we observed that BPA treatment for 24 hours did not change the total protein or K-48-linked ubiquitination profile (**Figure S1E and G**) but increased the K-63-linked ubiquitination (**Figure S1F**). However, 48 hours of BPA treatment did not alter any ubiquitination profile (**Figure S1E-G**). We performed the activity-based protein profiling of deubiquitinating enzymes, followed by western blot analysis, and found an altered enrichment of deubiquitinases between these conditions (**Figure S1H**), indicating a potential dysregulation of the ubiquitin-signalling network due to BPA exposure. Next, we treated the HTR8/SVneo cells with 10 μM BPA for 24 hours and performed activity-based deubiquitinase profiling using mass spectrometry under treated and control conditions, as illustrated in the schematic workflow in **Figure 2A**, to identify enriched active deubiquitinases (**File S1**). We identified a total of 19 deubiquitinases belonging to five deubiquitinase subfamilies, including ubiquitin C-terminal hydrolases (UCH) subfamily (UCHL1 and UCHL5), ubiquitin-specific proteases (USP/UBPs) subfamily (USP14, USP32, USP5, USP6, USP36, USP45, USP8, USP29, and USP17L1,2,5,8), ovarian tumor (OTU) domain family (OTUB1 and OTUD6), Machado-Joseph domain containing protease subfamily ( ATXN3), and Jab1/Pab1/MPN domain-containing (JAMM) protease subfamily (PSMD7 and PSMD14). Among the identified DUBs, 11 were found to be common in the BPA-treated and control conditions, 7 proteins were uniquely present in the control sample, and only one was in the treated sample (**Figure 2B**). The relative abundance of the identified deubiquitinases was represented as a heatmap of average emPAI values for control and treated samples (**Figure 2C**).

**Figure 2:**
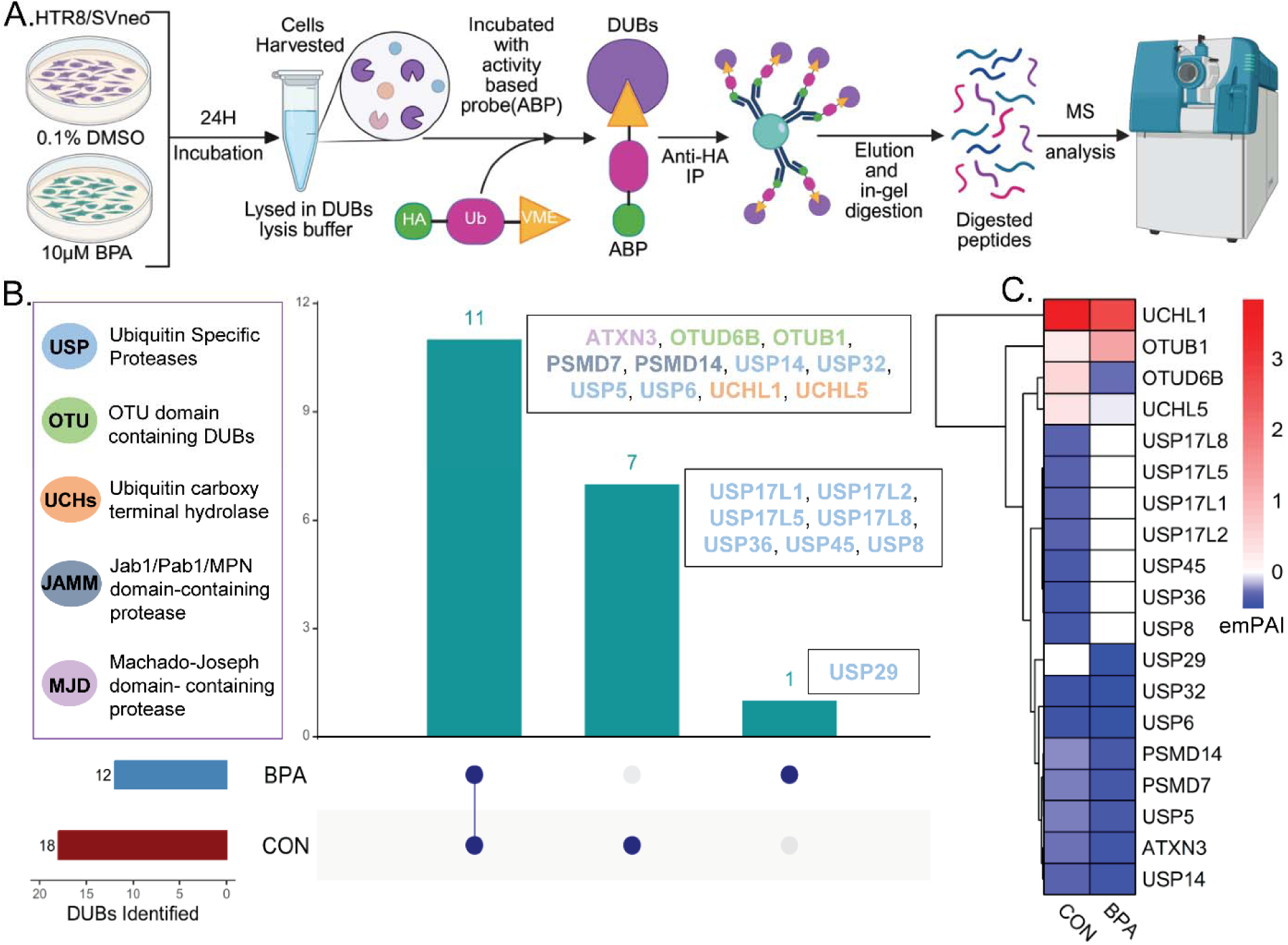
Active deubiquitinase profile alteration induced by low-dose BPA. **A)** Schematic workflow for enrichment of active deubiquitinases from BPA-treated and control samples using activity-based profiling probe HA-UbVMe. **B)** Upset plot showing the overlap of identified deubiquitinases between treated and control samples. The colour of the gene name mentioned on the right side of the bar indicates the subfamily associated with the deubiquitinases. The upper left panel of the plot indicates the deubiquitinase subfamily in colour-coded format. **C)** A heatmap showing the abundance of the identified deubiquitinases in treated or control samples. The gradient colour represents the average emPAI values for each identified deubiquitinase.

Previous studies have shown that BPA alters ubiquitin-signalling pathways [23], and our present findings corroborate this by demonstrating that BPA exposure regulates both ubiquitination events and deubiquitinase activity (**Figure 2A-D**). A few studies have demonstrated that the deubiquitinases like USP14 [41], USP8 [42], PSMD14 [43], USP5 [44], and UCHL5 [45] play a critical role in trophoblast function. The functions of many deubiquitinases in trophoblast cells or the placenta have not been explored. Moreover, none of the ubiquitin-signalling studies have been conducted in BPA exposure conditions. It is also important to note that the expression alteration of deubiquitinases affects the protein ubiquitination in cells, and these ubiquitinated proteins are the substrate for deubiquitinases. Therefore, we performed target substrate identification by performing global ubiquitinome analysis upon BPA treatment.

### Global ubiquitinated proteome map of extra-villous trophoblast cells

Protein quality control mediated by ubiquitination events in trophoblast cells is critical for the sustenance of a healthy placenta [24, 25]. Therefore, we performed ubiquitin-modified protein enrichment using the di-Gly antibody to examine the ubiquitination dynamics of HTR8/SVneo cells after BPA treatment. This approach involves immunoaffinity-based enrichment of trypsin-digested peptides modified with Gly-Gly at the ε-amino group of lysine through an iso-peptide bond, followed by comprehensive LC-MS/MS analysis of three bio-replicates from treated and control samples (**Figure 3A**). The assessment of MS data quality was done using the distribution of Posterior Error Probability (PEP) score and prot score for both modified and non-modified peptides and their associated proteins respectively. The mascot PEP score is statistical estimation of false discovery rate denoting confidence of peptide identification. A lower PEP score corresponds to more confidant identification of a peptide. The log2 distribution of the PEP score revealed more density at lower values for ubiquitinated peptides. Whereas, non-ubiquitinated peptides demonstrated a greater peak density at higher PEP score (**Figure 3B**). This suggests that ubiquitinated peptides were identified with greater confidence across all samples. Similarly, the mascot prot score, which is sum of the scores of the individual peptides matched to the protein filtered through stringent FDR, was plotted in log2 scale (**Figure 3C**). A higher prot score relates to better protein identification. Accordingly, our results showed that both the modified and non-modified proteins have considerably higher distribution of prot score, implying robust identification of ubiquitinated proteins with stringent modification filter. Therefore, by integrating stringent filtering criteria, specifically a reduced PEP score and elevated protein confidence scores, we identified a total of 794 ubiquitin-modified proteins and 3967 ubiquitinated peptides in at least one experimental condition (**File S2**). **Figure 3D** summarizes the individual identification of modified and unmodified peptides, as well as their associated protein groups.

**Figure 3:**
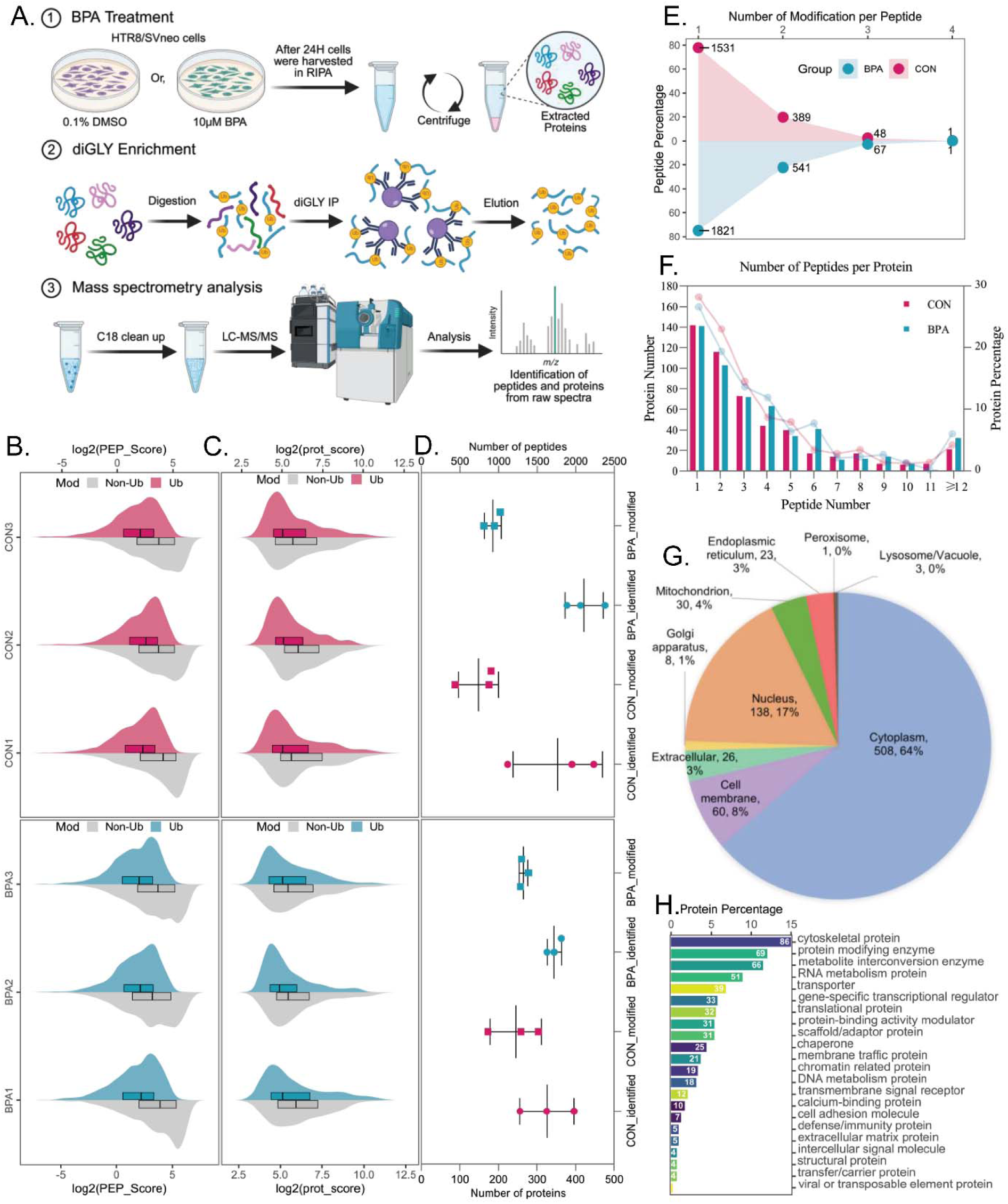
Ubiquitinome Landscape of trophoblast cells upon BPA exposure. **A)** Schematic workflow for global ubiquitinome profiling after BPA exposure. Gaussian distribution of **B)** PEP score for both ubiquitinated and non-ubiquitinated peptides and **C)** prot score for identified ubiquitinated and non-ubiquitinated proteins for all the biological replicates of treated and control samples. x-axis representing log2(PEP score) or log2(Prot score) respectively. **D)** Number of identified ubiquitin-modified and non-modified peptides (upper panel) and proteins (lower panel) in each biological replicate of BPA-treated and untreated samples represented as mean ± SEM. **E)** Distribution of multiplicity of modification on each peptide in BPA-treated and control samples. **F)** Count of peptides per identified proteins in BPA and control samples. **G)** Cellular localization of the Ub-modified proteins extracted from DeepLoc 2.0 using default parameters, represented by a pie-chart mentioning protein count and percentage for each cellular compartment. **H)** Fraction of ubiquitinated proteins identified based on protein functional classification using Panther (v19.0) webtool represented in a bar chart. x and y-axis represent protein percentage, and function, respectively. The number of proteins enriched for each function is denoted at the top of the bar.

After confirming a good data quality, we assessed the multiplicity of modifications on each peptide. Interestingly, we observed that many peptides were singly modified, followed by doubly and triply modified ubiquitinated sites. BPA-treated samples exhibited a higher number of modification sites, irrespective of whether peptides carried single or multiple modifications (**Figure 3E**). The peptide count distribution for each ubiquitinated proteins demonstrated the identification more than 20% of total reported protein with single peptides. The protein number gradually decreases with increasing peptide counts observed for a protein. The pattern remains consistent for treated and control samples (**Figure 3F**). The sub-cellular localization analysis of ubiquitinated proteins showed that the majority of protein are from cytoplasmic fraction (64%) but rest 36% proteins are distributed evenly among other cellular compartments (**Figure 3G**). Protein functional class prediction using Panther (v19.0) database majorly enriched cytoskeletal proteins (N=86) and protein-modifying enzymes (N=69), both associated with migration and ubiquitin modification events, strengthening our initial findings in this study (**Figure 3H**).

### BPA alters the ubiquitination landscape of the trophoblast cells

Next, we explored the ubiquitinome landscape of HTR8/SVneo cells and identified a dynamic pool of ubiquitin-modified proteins. We inspected the uniquely modified proteins present in the BPA-exposed group to reveal the BPA-induced alteration. We observed that the BPA-treated and control groups have 280 and 250 unique ubiquitinated proteins, respectively, and 264 ubiquitinated proteins common to both groups (**Figure 4A**). To delineate the BPA-specific alteration, we chose the 280 unique proteins and verified their functional association using protein-protein interaction (PPI) network and pathway enrichment in Metascape (v3.5.20250701). The physical interaction network of unique ubiquitinated proteins compiled using STRING and BioGrid, which provided a comprehensive interaction map. This map highlighted a densely connected network of clusters, which was achieved using the Molecular Complex Detection (MCODE) algorithm (**Figure 4B**). The algorithm detected 9 different MCODE clusters annotated using multiple colours. Further, pathway enrichment of these PPI clusters described their functional representation. The best three pathways by p-value were used to represent the functional description of the PPI cluster (**Figure 4C**). We observed two PPI clusters, like MCODE2 and MCODE9, enriched pathways, such as intermediate filament cytoskeletal organization, microtubule-based transport, and movement, respectively. The proteins involved in this two PPI cluster are KRT20, KRT40, KRT23, KRT33B, KRT33A, VIM, TPMO, DDX3X, VCL, MYH13, MYL6, MYL9, NIN, VCL in MCODE2 and IFT81, TRAF3IP1, KATNIP in MCODE9 cluster. This finding highlighted the association of altered migration events with a unique ubiquitinated proteome altered by BPA. Other significant MCODE clusters enriched functional categories like the complement system in neuronal development (MCODE1), cytoplasmic translation (MCODE3), Activation of AMPK downstream of NMDARs (MCODE4), mRNA processing (MCODE5), Kinesins (MCODE6), C complex spliceosome (MCODE7), and PID RHOA REG pathway (MCODE8) (**Figure 4C and File S3**). Pathways enriched from PPI clusters provided some snapshot of BPA-mediated dysregulation of the ubiquitinated proteome. To get a better perspective, we explored the network of terms enriched from all the uniquely modified proteins in the BPA group. The network was built using hierarchical clustering (Kappa score *>*0.3) of pathways and processes enriched with a p-value *<*0.01, a minimum gene count of 3, and an enrichment factor *>*1.5. Each cluster was represented by its most statistically significant term, along with protein count as bubble size and q-values for significance within the clusters (**Figure 4D**). The network enriched multiple migration-related pathway clusters, such as motor proteins, microtubule-based movement, microtubule cytoskeleton organization, intermediate filament organization, and plasma membrane-bound cell projection assembly, with the most significant q-values and high protein counts for each cluster term (**Figure 4D and File S4**). Other interesting pathways relate to metabolism of RNA, mitotic cell cycle, positive regulation of translation in response to ER stress, mitochondria localization and oncogenic MAPK signalling.

**Figure 4:**
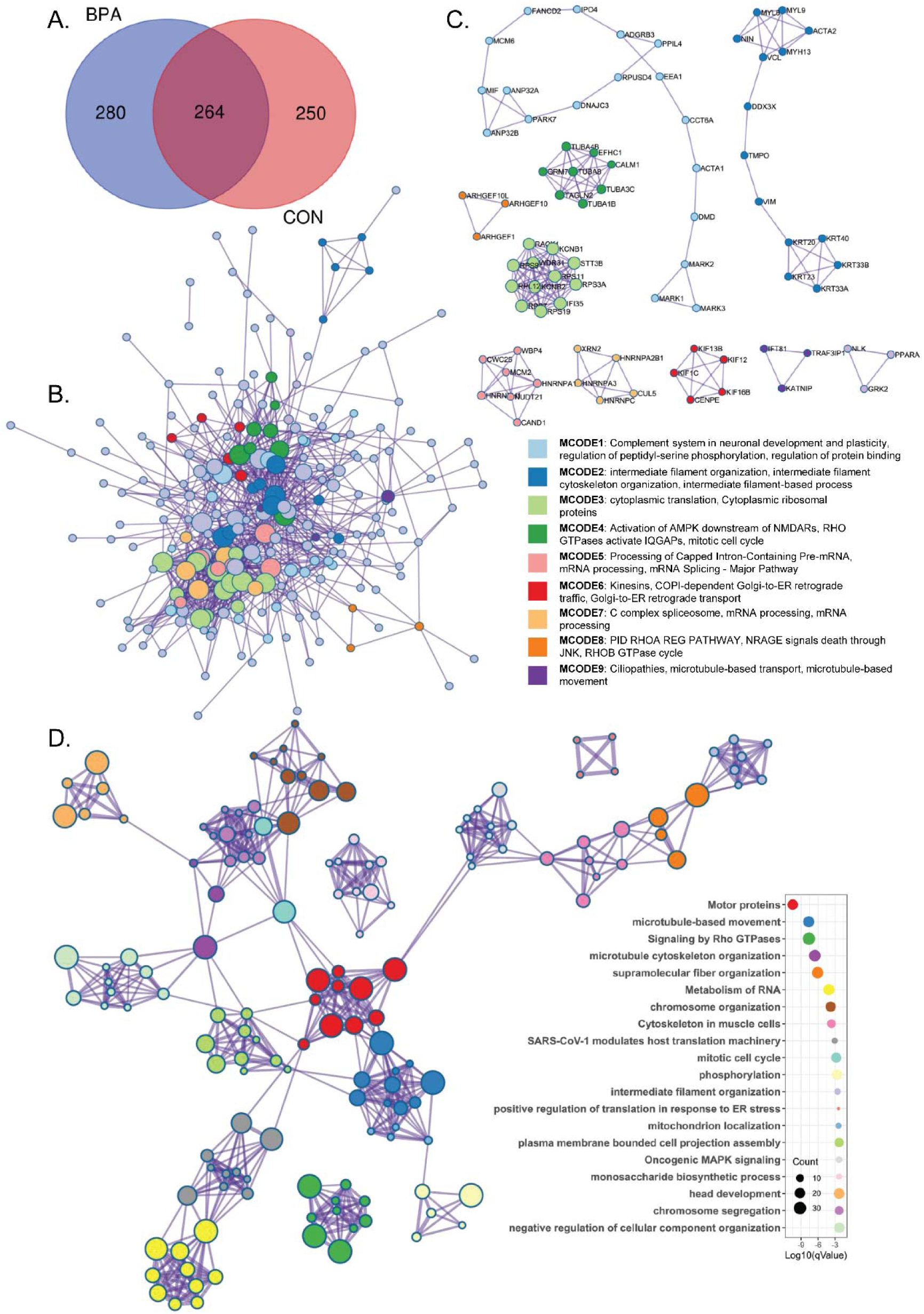
BPA triggers alteration of ubiquitination dynamics in HTR8/SVneo cells. **A)** Venn diagram representing common and unique ubiquitinated proteins between BPA exposed and control condition. **B)** Protein-protein interaction (PPI) network of BPA altered ubiquitinylated proteins using only physical interactions from STRING (physical interaction score *>*0.132) and BioGrid. Each circle represents a protein and edge length represents physical interaction score between a pair. The Molecular Complex Detection (MCODE) algorithm is employed to identify densely connected network component. The colour of each circle reflects its associated MCODE network. **C)** PPI cluster of associated proteins from each coloured MCODE component. Three best scoring pathways or process terms based on p-value is retained as functional description of each MCODE component. **D)** A network of enriched pathways due to BPA exposure was plotted using terms with connected edges having similarity *>*0.3. The node colour demonstrates the cluster identity, shown as in a bubble plot, with the x-axis plotted as log10(Q-value) and the size of the bubble denoting the count of proteins for the respective cluster.

Similar analysis using ubiquitinylated proteins in control groups yielded cluster terms like signalling by Rho GTPase, cell cycle, metabolism and localization of RNA, chromosome organization and remodelling, glucose-6-phosphate, monosaccharide and DNA metabolic processes (**Figure S2 and File S4**). These cluster terms and their associated ubiquitinated proteins are essential for the sustenance of healthy HTR8/SVneo cells. An identical PPI enrichment analysis of unique proteins from control samples fetched five MCODE PPI clusters which are metabolism of RNA (MCODE1 & 2), metabolic reprogramming in colon cancer (MCODE 3), rRNA processing in the nucleus and cytosol (MCODE4), and COPI-dependent Golgi-to-ER retrograde traffic (MCODE5) (**Figure S2C-D and File S3**). None of the MCODE clusters or pathway networks from control group overlaps with migration related PPI cluster or network term enriched in BPA group suggesting the alteration of migration-related protein ubiquitination was very specific to BPA-exposed state. Although, the altered ubiquitinome shows directional regulation of migration related phenotype, the underlying molecular mechanism is not well understood. Therefore, we integrated the previous DUBs profiling information with the BPA-exposed ubiquitinome signals to unravel the molecular players of BPA-induced migration alteration.

### Integrated substrate-ligase-deubiquitinase network highlights key nodes altered by BPA

The protein homeostasis is maintained by tight regulation of deubiquitinase and E3 ubiquitin ligase expression and activity [24]. Hence, dysregulation of deubiquitinases and the E3 ubiquitin ligase may be responsible for the altered ubiquitination profile in BPA-exposed conditions. In this study, we identified 19 deubiquitinases through activity-based DUB profiling. To evaluate their functional relevance, we integrated substrate information and extracted 2,056 potential substrate targets from Ubibrowser 2.0. We compared the Ubibrowser-predicted proteins with our identified ubiquitinated proteome and identified 33 common substrates that were unique to the BPA-treated group (**Figure 5A**). These 33 common substrates were covered by only 10 deubiquitinases from the list of 19 DUBs identified. We performed Reactome pathway enrichment of these 33 proteins and observed caspase-mediated cleavage of cytoskeletal proteins with the highest fold enrichment supported by other key pathways such as NRAGE signals death through JNK, apoptotic cleavage of cellular proteins, Rho GTPase cycle, etc (**Figure 5B and File S5**). To integrate the substrate-ligase-deubiquitinase network, we again enriched 61 known and putative E3 ligases for 33 substrates with a confidence score *>*0.8 from the Ubibrowser 2.0 database. This network of 61 putative or known ligases (violet circle), 10 deubiquitinases (green hexagon), and 33 ubiquitinated substrates (blue triangle) was connected using Ubibrowser confidence score as edge width and visualized in Cytoscape (**Figure 5C and File S5**).

**Figure 5:**
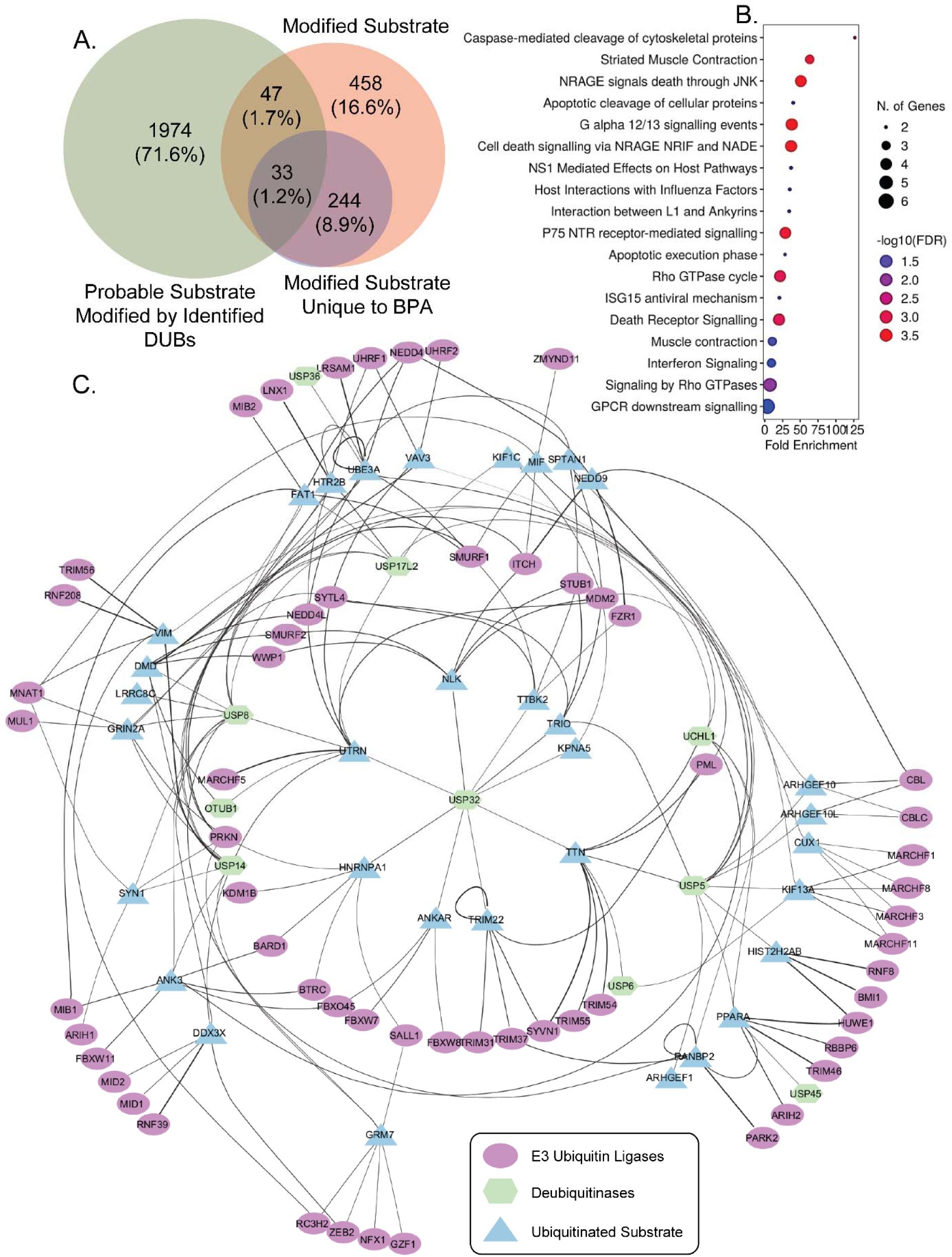
Integrated ubiquitinated substrate, their target deubiquitinase, and E3 ubiquitin ligase network after BPA exposure. **A)** Venn diagram showing overlap between probable substrate target for identified deubiquitinase, all modified substrate identified in this study, and modified substrate unique to only BPA group represented as proteins count and its percentage. **B)** Reactome pathway enriched by the 33 common ubiquitinated substrates illustrated using a bubble plot, where the x-axis shows fold enrichment, while bubble color and size correspond to, log10(FDR) and protein count, respectively. **C)** The integrated network of 33 common substrates, their probable deubiquitinase (N=10) identified in DUBs profiling, as well as their known and probable E3 ubiquitin ligase (N=61) with a confidence score *>*0.8. The edge width denotes the confidence score extracted from Ubibrowser 2.0, and the network is visualized using Cytoscape. The violet circle represents the E3 ubiquitin ligase, the green hexagon is the deubiquitinases, whereas the blue triangle is the ubiquitinated substrate.

Next, we investigated the association of BPA with the 33 common substrate by direct or indirect means. We find out mRNA level dysregulation of Homeobox protein cut-like 1 (CUX1) [46] and Glutamate receptor ionotropic 2A (GRIN2A) [47] are reported in mouse brain hippocampus due to prenatal exposure of BPA. High dose of BPA administration also disrupted the DNA methylation at the ubiquitin-protein ligase E3A (UBE3A) locus, resulting in reduced UBE3A mRNA levels in both the placenta and brain tissue of mice [48]. Male mice prenatally exposed with BPA shows increased expression of the presynaptic marker Synapsin-1 (Syn1) in primary hippocampal neurons [49]. Further inspection shows, BPA stimulated the release of Macrophage migration inhibitory factor (MIF), a pro-inflammatory cytokine, from human endometrial [50] and decidual cells [51]. BPA also downregulates the epithelial to mesenchymal transition mediator vimentin (VIM) in HTR8/SVneo cells [18]. Interestingly, it has been reported that the BPS, a BPA analogue, triggers Peroxisome proliferator-activated receptor alpha (PPAR-alpha), a ligand activated trsacription factor and mediates the transcriptional activation of downstream genes in hepatocytes [52]. BPA and its other analogues also promote PPAR-alpha induced testicular dysfunction [53]. PPAR-alpha, along with its isoforms PPAR-gamma and PPAR-delta, is reported to bind BPA and disrupt its target genes’ regulation [27,54,55]. Moreover, increased activation and nuclear translocation of PPAR-alpha led to reduced expression of MMP2 and MMP9, along with impaired cellular migration in HTR8/SVneo cells [56]. These findings allowed us to correlate BPA-induced activation of PPAR-alpha with the reduced migration phenotype earlier found in this study.

### BPA triggers hyper-ubiquitination of PPAR-alpha in trophoblast cells

Peroxisome proliferator-activated receptors are highly expressed in trophoblast cells and are essential for various physiological functions like migration and invasion [57]. PPAR-alpha specifically binds with co-activators like fibrates or fatty acids, facilitating the formation of a complex with retinoic X receptor alpha, which then recognizes DNA sequences called peroxisome proliferator responsive elements (PPREs) to promote the expression of genes associated with lipid metabolism, energy balance, and inflammatory response [58]. The ubiquitination of PPAR-alpha affects its stability and signalling pathways, leading to altered expression of its target genes [59]. Therefore, to validate BPA-mediated hyper-ubiquitination of PPAR-alpha, we pulled PPAR-alpha from control or BPA-treated lysate and probed with anti-ubiquitin as well as anti-PPAR-alpha antibody. We observed a substantial increase in the mono-ubiquitinated fraction only in the BPA-treated lane when developed against the anti-ubiquitin antibody. Interestingly, the anti-PPAR-alpha antibody showed two bands, possibly modified and unmodified PPAR-alpha, in pulldown samples. However, the upper band in the BPA-treated condition shows increased intensity (**Figure 6A**). This result implies that BPA exposure indeed promotes the ubiquitination of PPAR-alpha in HTR8/SVneo cells.

**Figure 6:**
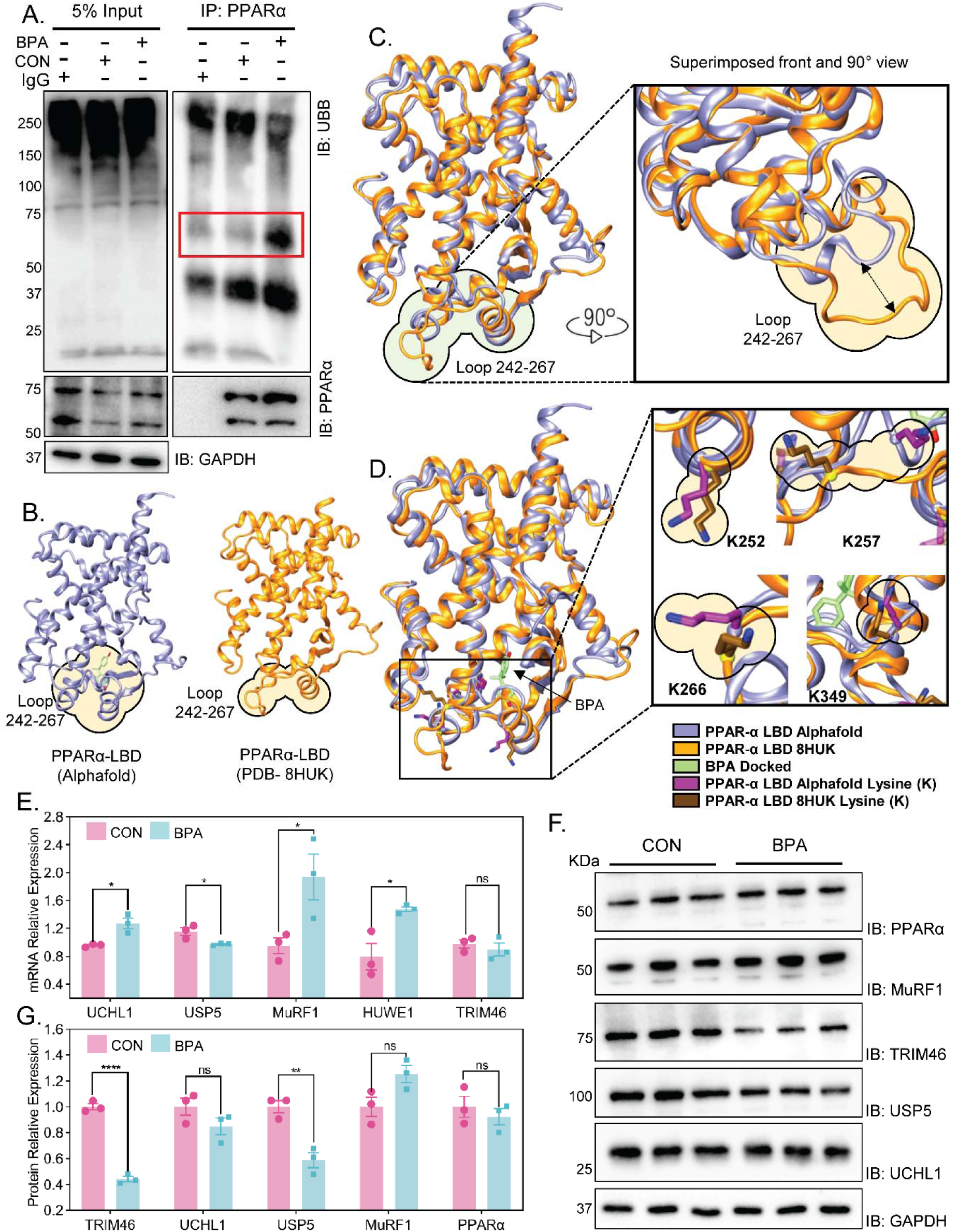
BPA mediates hyper-ubiquitination of PPAR-alpha. **A)** Pull-down of PPAR-alpha is performed to assess its ubiquitination profile after BPA administration. PPAR-alpha antibody and its isotype control IgG antibody were used for pulling PPAR-alpha from the protein lysate. The PPAR-alpha enriched samples were immunoblotted against anti-ubiquitin and anti-PPAR-alpha antibody, along with the input samples from un-enriched fractions. The red highlighted region indicates the mono-ubiquitination of PPAR-alpha in BPA samples. **B**) Individual structures of PPAR-alpha-LBD from the AlphaFold model (ice-blue) and the crystal structure PDB 8HUK (orange) are shown, with the 242–267 loop region circled. The AlphaFold model represents the dominant conformation observed in most ligand-bound PPAR-alpha crystal structures, whereas the 8HUK structure displays the loop in an open conformation. (**C**) Superimposition of the AlphaFold and 8HUK structures reveals positional shifts of the 242-267 loop. The docked pose of BPA (green) and the key lysine residues K252, K257, K266, and K349 are highlighted to illustrate their spatial relationship to the ligand-binding pocket. (**D**) Close-up views of the lysine side chains show their orientation within the dynamic loop region and proximity to the BPA binding site. **E**) Transcript abundance of UCHL1, USP5, MuRF1, HUWE1, and TRIM46 was measured utilizing qRT-PCR and normalized to GAPDH expression using the ΔΔCt method. Results are represented as mean ± SEM from three independent biological replicates. Multiple students’ t-test was employed, considering p-value < 0.05 as significant. **F)** Western blot analysis of PPAR-alpha, MuRF1, TRIM46, USP5, and UCHL1 from three biological replicates of BPA-exposed and control samples. GAPDH is used as a loading control for the respective experiments. **G)** The densitometric analysis of acquired immunoblots is summarized as mean ± SEM (N=3). The statistical significance was evaluated using multiple Student’s t-tests. The differences were considered significant at *p* < 0.05.

Although the mode of increased PPAR-alpha ubiquitination can have two proposed mechanisms. Firstly, the direct binding of BPA to PPAR-alpha may drive a structural change in conformation, exposing the identified residues of K252, K257, K266, and K349 (**Figure S3A**), making them accessible for the E3 Ubiquitin ligases, as seen in the case of PPAR-gamma [27]. To explore this mechanism, we performed molecular docking studies, which revealed that BPA occupies the same binding region in PPAR-alpha (alpha-fold) as it was previously reported in the co-crystal structure of PPAR-gamma, with no steric clashes. However, the binding pose of BPA at PPAR-alpha is slightly different than PPAR-gamma, suggesting some residual changes in conformation that prevent BPA from achieving a similar orientation (**Figure S3B-E**). Since we are exploring BPA binding and its association with the ubiquitination process, we compared the structural changes where key lysine residues (K252, K257, K266, and K349) are localized (loop spanning residues 242-267) using alpha-fold and reported crystals. It is also interesting that this loop is part of the BPA binding site as well, indicating the possibility of this region being involved in the association of BPA binding and facilitating ubiquitination, which was observed in the BPA-exposed ubiquitinome.

To gain mechanistic insight as to how BPA binding is involved and its possible association with ubiquitination, we explored the BPA binding site and conformational orientation of lysine residues lying in the loop. We manually inspected all 69 PPAR-alpha crystal structures deposited in the RCSB PDB, mainly focusing on the loop. This analysis revealed that the loop spanning residues 242-267, which contains all aforementioned lysine residues except K349, exhibits substantial conformational plasticity with maximum positional shifts of up to ∼15.0 Å across different structures. Distance measured on the superimposed structure of the bound and unbound loop of PPAR-alpha by taking residue lysine 257 (**Figure S3F**). Such structural variability indicates a highly dynamic loop capable of sampling multiple conformations. Structural evidence further indicates that this loop can form a helical conformation upon ligand binding, as observed from mining different reported cocrystals. Among the examined structures, PDB ID: 8HUK [60] was particularly informative, as it contains a biological assembly lacking a bound ligand and displays this loop in an open conformation, further supporting its flexibility in the absence of a ligand (**Figure 6B-D**).

To quantitatively evaluate the stability of this loop, we performed normal mode analysis (NMA) using the ProDy plugin of VMD. The NMA results revealed that BPA binding significantly increases the stability of the 242-267 loop region (∼10-fold) and the lysine residues embedded within it (**Figure S3G-H**). This enhanced stabilization may promote more favourable interactions with ubiquitin, thereby facilitating ubiquitination at these lysine sites. These findings suggest that BPA not only occupies the PPAR-alpha binding pocket but also enhances the stability of the dynamic regulatory loop and lysine residue, which may promote favourable interactions with ubiquitin.

To cross-check these observations, as ubiquitin binding occurs through the formation of an iso-peptide bond between the terminal glycine of ubiquitin and the lysine of the target protein, we conducted protein-protein docking studies with bound (with BPA) and unbound open form of PPAR-alpha (8HUK). In the BPA unbound open form, the docked poses of ubiquitin are mostly far away from loop regions, indicating the loop region is less preferable. However, in the BPA-bound form, we observed ubiquitin poses localized near loop regions. From the analysis, we also observed that the terminal glycine of the ubiquitin docked pose is also able to form iso-peptide bonds, as the measured distance is 2.3 Å with K257. While in unbound form, the distance between them is 12.8 Å (ubiquitin@G76: PPAR-alpha: K257). This analysis seems true for other lysines as well (**Figure S3I-J**), indicating direct evidence that BPA-bound states are more stable. Overall, these findings indicate that BPA binding seems to promote a conformational state of PPAR-alpha that facilitates ubiquitin recognition and supports ubiquitination. Importantly, the structural stabilization comes from the ligand-binding pocket itself; it is likely that other ligands binding to the same site could cause similar effects to stabilize the flexible 242-267 loop and influence ubiquitination in a similar fashion.

The alternative mechanism could also involve the indirect regulation of known or predicted deubiquitinase or E3 ubiquitin ligase of PPAR-alpha after BPA exposure. Thus, we investigated the substrate-Ligase-deubiquitinase network (**Figure 5C**) and highlighted the PPAR-alpha node. The PPAR-alpha node relates to 3 deubiquitinases, namely UCHL1, USP5, and USP45, and five E3 ligases including HUWE1, TRIM46, RBBP6, ARIH2, and RANBP2 (**Figure S4A**). It has been reported earlier that HUWE1, TRIM46, and MuRF1 participate in ubiquitination of PPAR-alpha. Interestingly, HUWE1 polyubiquitinates PPAR-alpha in hepatocytes [61]. On the other hand, TRIM46 performs the PPAR-alpha’s ubiquitination in osteosarcoma [62]. A separate study demonstrates that MuRF1, a muscle-specific E3 ligase, mono-ubiquitinates PPAR-alpha, facilitating its nuclear export in cardiomyocytes and inhibiting fatty acid oxidation through a proteasome-independent process [63]. However, no DUBs have been identified to target the de-ubiquitination of PPAR-alpha in any context. Thus, we curated a list of three known Ligases and two probable DUBs of PPAR-alpha for qRT-PCR and western blot analysis.

Interestingly, we observed a significant RNA level up-regulation of UCHL1, MuRF1 and HUWE1 after BPA treatment. However, USP5 and TRIM46 showed a down-regulation trend but not very significant (**Figure 6E**). Despite finding a significant increase in HUWE1 expression at the RNA level, we did not evaluate the protein expression, as HUWE1 mediates PPAR-alpha poly-ubiquitination, and we observed mono-ubiquitination of PPAR-alpha in response to BPA exposure (**Figure 6A**). Next, we observed that MuRF1 protein is significantly upregulated with a 1.3-fold change. Whereas TRIM46 and USP5 showed significant protein level down-regulation. The protein expression of PPAR-alpha and UCHL1 remained unchanged (**Figure 6D and F**). These results indicate that BPA exposure enhances MuRF1 expression, which then mono-ubiquitinates PPAR-alpha. In contrast, BPA down-regulates USP5, leading to a reduction in PPAR-alpha de-ubiquitination. This reciprocal regulation of MuRF1 and USP5 due to BPA administration results in the accumulation of mono-ubiquitinated PPAR-alpha in trophoblast cells. Our findings imply that both proposed mechanisms may contribute to the buildup of mono-ubiquitinated PPAR-alpha in BPA-exposed trophoblast cells. However, the underlying mechanism by which the mono-ubiquitinated PPAR-alpha influences the decreased migration phenotype post-BPA treatment in trophoblasts is yet to be elucidated, which encouraged us to explore the fate of the mono-ubiquitinated PPAR-alpha in HTR8/SVneo cells.

### Nuclear localization of ubiquitinated PPAR-alpha alters migration-related gene expression

PPAR-alpha is a ligand-responsive transcription factor, and its nuclear translocation is critical for the regulation of its transcriptional activity. PPAR-alpha harbors at least two nuclear localization signals in the DNA-binding domain (DBD)-hinge and activation function 1 (AF1) region, which facilitates its nuclear shuttling [64, 65]. MuRF1 is known to mono-ubiquitinate PPAR-alpha, enhancing its nuclear export and inhibiting its target gene regulation [63]. These findings guided us to investigate the localization of mono-ubiquitinated PPAR-alpha in HTR8/SVneo cells post-BPA treatment. The immunostaining of the BPA-exposed and control cells with anti-ubiquitin (Red channel) and anti-PPAR-alpha (Green channel) antibodies to visualize co-localization of these two proteins inside the cell, specifically in the nucleus (**Figure 7A**). We observed enhanced localization of ubiquitinated PPAR-alpha in the nucleus after BPA treatment (**Figure S4D**). This nucleus-localized ubiquitinated PPAR-alpha can repress or induce its target gene expression.

**Figure 7:**
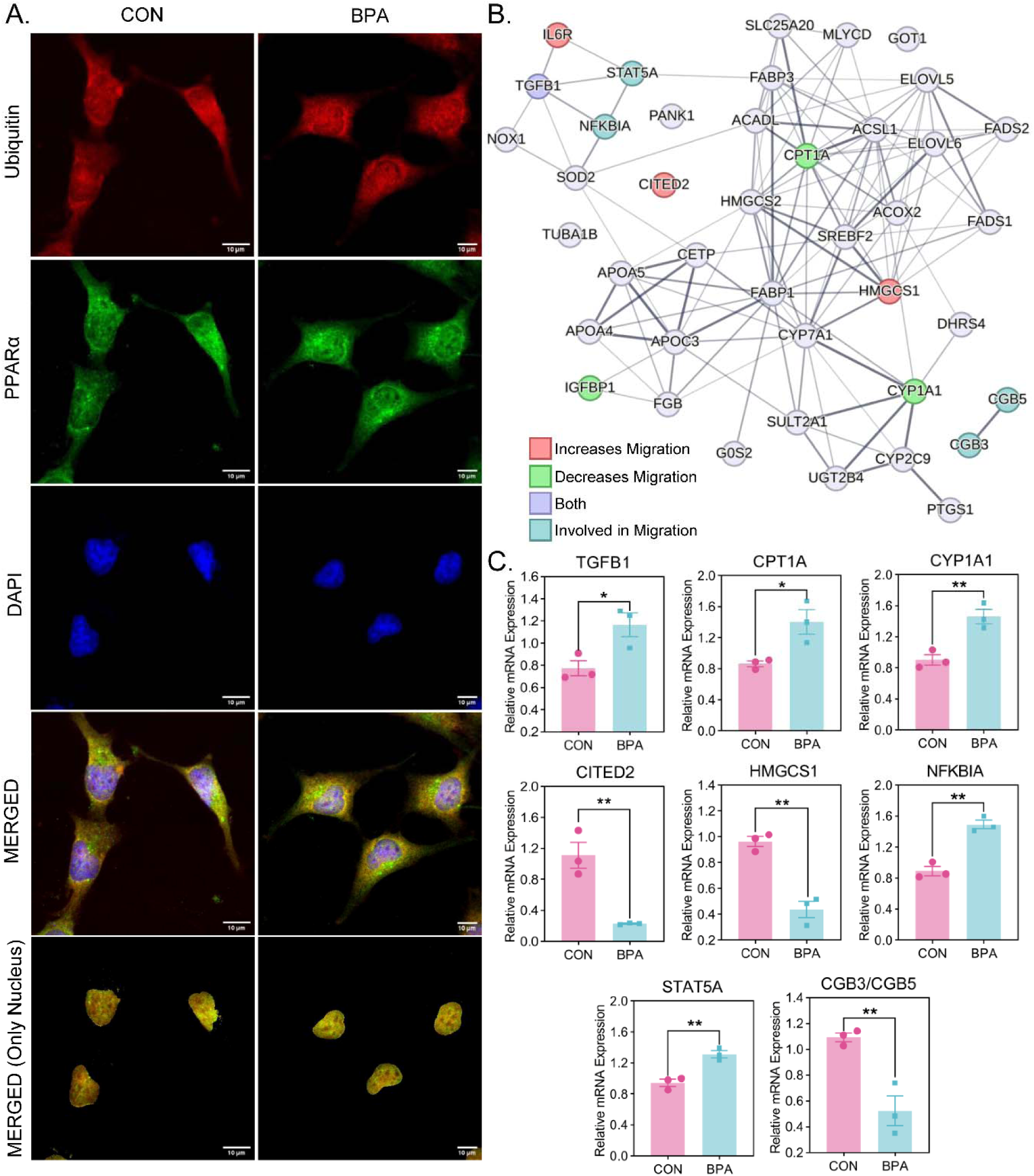
Ubiquitination enhances nuclear retention of PPAR-alpha, disrupting its target gene expression. **A)** Subcellular distribution of ubiquitinated PPAR-alpha in BPA-treated HTR8/SVneo cells visualized using confocal microscopy. Cells were immune-stained with anti-PPAR-alpha (green) and anti-ubiquitin (red) antibodies. While for the nucleus, the cells were counter-stained using DAPI (blue). Under basal conditions, ubiquitinated PPAR-alpha localizes primarily in the cytosol, and upon BPA exposure, it translocates to the nucleus. Images were captured using a Leica SP8 confocal microscope with a 63X oil objective. Scale bars = 10 μm. **B)** STRING-enriched protein-protein interaction network of 41 unique target genes of PPAR-alpha highlights the proteins involved in migration, curated from literature. Node colours demonstrate the association of genes with different migration phenotypes mentioned below the network. **C)** qRT-PCR was performed using gene-specific primers for TGFβ1, CPT1A, CYP1A1, CITED2, HMGCS1, NFKBIA, STAT5A, and CGB3/CGB5 and normalized to GAPDH utilizing the ΔΔCt method. Each data points reflects the mean ± standard error of the mean (SEM) from three independent biological replicates. Statistical analysis was performed using Student’s t-test and significant differences were considered at *p* < 0.05.

Previous findings suggested that nuclear localization of PPAR-alpha leads to reduced migration and invasion in HTR8/SVneo cells [56]. In this study, we found that BPA treatment caused reduced migration and elevated nuclear localization of ubiquitinated PPAR-alpha. Therefore, to link these two observations, we fetched the genes transcriptionally targeted by PPAR-alpha along with its other two isoforms from the TFLink database. Next, we overlapped only the validated targets of the three PPAR isoforms (PPAR-alpha, PPAR-beta, and PPAR-gamma) to isolate 41 targets unique to PPAR-alpha only (**Figure S4B and File S6**). The KEGG pathway enrichment of these 41 unique PPAR-alpha targets revealed metabolism of Fatty acid, cholesterol, and biosynthesis of unsaturated fatty acid as the most significantly enriched pathways (**Figure S4C and File S6**).

PPAR-alpha is already known to regulate lipid metabolism genes [66]. Hence, to identify migration-associated genes from the list of 41 enriched targets, we performed a literature survey. We found that Transforming growth factor-β 1 (TGF-β1) can stimulate [67] or repress [68] HTR8/SVneo cells migration in a context-dependent manner. However, genes like Carnitine palmitoyl transferase 1A (CPT1A) [69], cytochrome P450 monooxygenase 1A1 (CYP1A1) [70] and Insulin-like growth factor-binding protein 1 (IGFBP1) [71] inhibit migration in HTR8/SVneo cells. Similarly, Glutamic acid/aspartic acid-rich carboxyl-terminal domain 2 (CITED2) [72], Hydroxy-methyl-glutaryl-CoA synthase (HMGCS1) [73] and Interleukin-6 receptor (IL-6R) [74] are associated with elevated migration in HTR8/SVneo cells. This key migration regulator, along with other migratory signaling-associated genes such as NF-kappa-B inhibitor alpha (NFKBIA), signal transduction and activation of transcription 5A (STAT5A), and Beta subunit of the human chorionic gonadotropin (CBG3/5), are highlighted in the STRING network of those 41 unique PPAR-alpha targets. The genes are annotated with colour according to their associated role in migration regulation (**Figure 7B**). Finally, we performed qRT-PCR of these shortlisted genes and found significant up-regulation of TGFβ 1, CPT1A, CYP1A1, NFKBIA, and STAT5A upon BPA exposure (**Figure 7C**). Whereas CITED2, HMGCS1, and CGB3/5 show significant downregulation (**Figure 7C**). In line with our literature survey, we found genes associated with migration inhibition (CPT1A & CYP1A1) are up-regulated post-BPA treatment, but genes linked with increased migration (CITED2 & HMGCS1) are down-regulated due to BPA. We also reported the regulation of NFKBIA, STAT5A, and CBG3/5 proteins, which were involved in the regulation of cellular migration in a context-specific manner. Although IGFBP1 and IL-6R show regulation opposite to the literature-suggested direction, indicating an alternative modulation of these proteins by BPA (**Figure S4C**).

## Discussion

This study demonstrates that low-dose BPA disrupts trophoblast cell migration without inducing apoptosis, indicating a BPA induced functional dysregulation rather than cytotoxic effect. The observed migration impairment correlates with reduced MMP2 and MMP9 expression and aligns with previous reports of BPA mediated trophoblast dysfunction. Migration requires precise regulation of protein turnover, and our data reveal that BPA selectively remodels the ubiquitin-proteasome signaling network. Activity-based profiling of deubiquitinases in trophoblast cells identified 19 deubiquitinating enzymes. This deubiquitinases shows a coordinated pattern of dysregulation upon BPA administration. Global ubiquitinome profiling uncovered a BPA induced unique ubiquitination signature in trophoblast cells. This altered ubiquitinated protein pool enriched pathways involving cytoskeletal organization, motor protein function, and cell projection assembly. The absence of these signatures in control cells underscores the specificity of BPA-induced ubiquitin signaling alterations. Integrated network analysis identified PPAR-alpha as a one of the major regulatory node affected by BPA. Molecular dynamics stimulation reveals a novel mechanism by which BPA directly binds to PPAR-alpha, stabilizing a flexible lysine-rich loop and exposing K252, K257, K266, and K349 residues, thereby promoting mono-ubiquitination of PPAR-alpha. The down regulation of deubiquitinase USP5 and upregulation of E3 Ligase MuRF1 upon BPA treatment may also elevate the ubiquitination of PPAR-alpha. Both mechanisms may increase the ubiquitinated pool of PPAR-alpha in cytosol, triggering its nuclear translocation. In the nucleus, it up-regulates migration-inhibiting gene expression (CPT1A, CYP1A1) and down-regulates genes (CITED2, HMGCS1) required for cellular migration. This cumulative effect reduces the migration of HTR8/SVneo cells upon BPA administration and disrupts their physiological functions (**Figure 8**). We assume this molecular-level dysregulation by BPA will also affect the placental homeostasis, which may contribute to placental dysfunction associated with pregnancy complications. Overall, our findings establish a previously unrecognized BPA-PPAR-alpha-ubiquitin axis that disrupts trophoblast migration through coordinated structural, post-translational, and transcriptional regulation. These insights provide mechanistic depth to the adverse effects of environmental BPA exposure on placental function and pregnancy outcomes. While the results are encouraging, the present study still has several limitations. Initially, the activity based deubiquitinase profiling only captured the active deubiquitinases in trophoblast. while some deubiquitinase with non-canonical signalling may also be affected by BPA. Also, this current study is performed in an isolated cell culture model not considering how BPA can regulate the protein homeostasis in tissue level. Subsequent mice studies will be helpful to understand the effect of BPA on systems level during pregnancy. Finaly, this study considered that chronic exposure to BPA during pregnancy is one of the key factors for placental dysfunction in adverse pregnancy outcomes. Although, the extent of BPA exposure during pregnancy is not measured in this study which reduces the scope of comparing the clinical outcome with BPA exposure.

**Figure 8:**
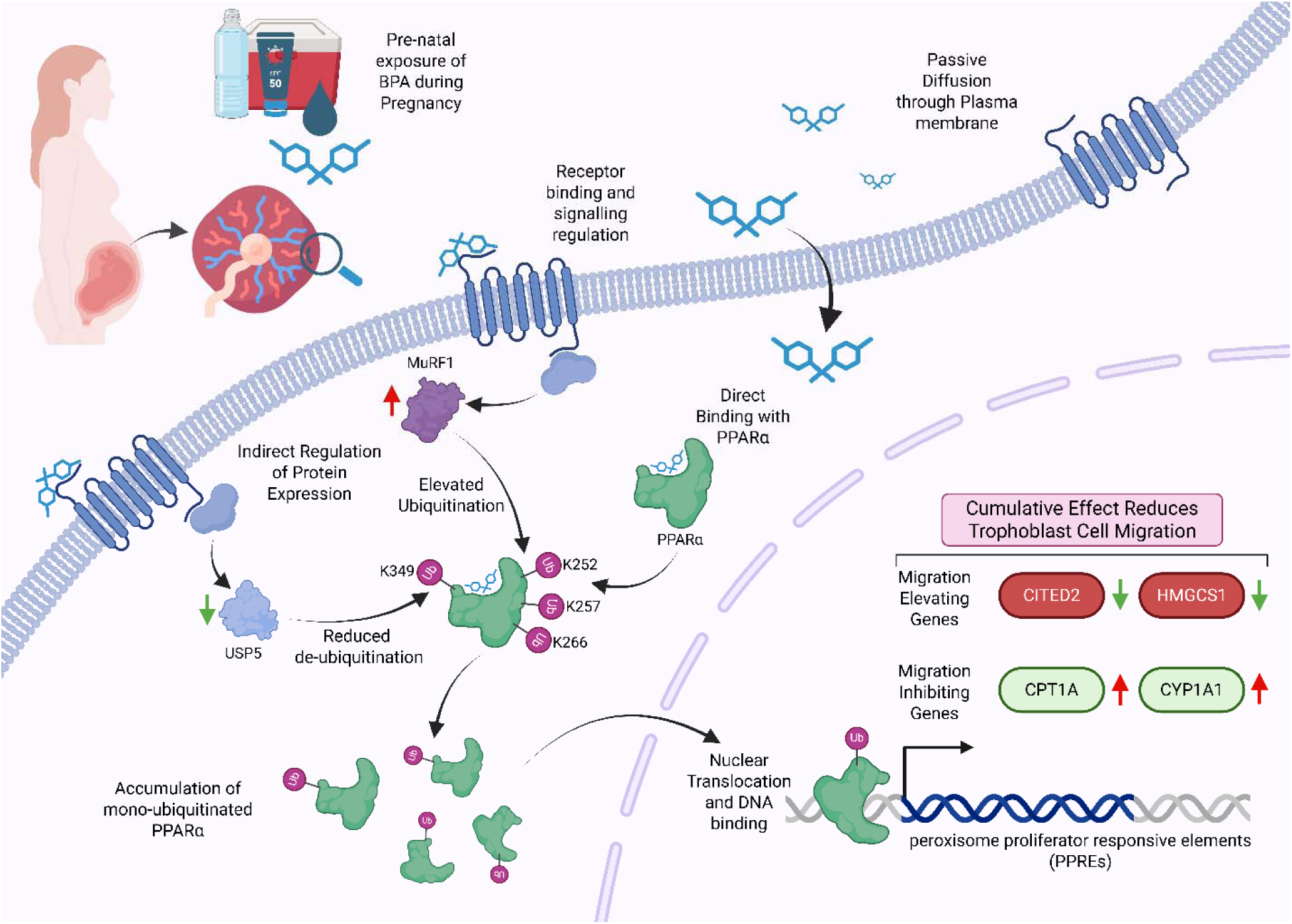
Graphical representation summarizing the key findings. The image depicts mechanistic insights of BPA-mediated PPAR-alpha ubiquitination via direct BPA binding, which stabilizes the K252, K257, and K266 residues or indirect regulation of E3 ligase MuRF1 and DUBs USP5. Both facilitate the build-up of mono-ubiquitinated PPAR-alpha upon BPA exposure. This ubiquitinated PPAR-alpha localizes to the nucleus and dysregulates key migration-associated target genes, which ultimately contribute towards reduced migration phenotype in HTR8/SVneo cells.

## Author contributions

T.K.M. and A.B. conceived the project and designed the experiments. A.B. and S.S. performed proteomics sample preparation, mass spectrometry data acquisition, and analysis. A.B. performed, and K.S.B. helped in biochemical and cellular experiments. D.S. and S.A. helped in molecular docking studies. A.B. and T.K.M. wrote the manuscript. All authors read and approve the final version of the manuscript.

## Acknowledgements

TKM acknowledges the Regional Centre for Biotechnology for its intramural research funding. We thank Dr. Pallavi Kshetrapal for providing us with the HTR8/SVneo cell line. We express our sincere gratitude to all the technical staff at RCB for keeping the equipment in functional condition. We are grateful to all the members of the laboratory of functional proteomics for maintaining an excellent research environment. Debapriyo Sarmadhikari thanked THSTI, Sandhini Saha thanked ICMR, Ankit Biswas thanked DBT and Krishna Singh Bisht thanked DST-INSPIRE for their fellowship.

## Data and code Availability

The mass spectrometry data are available online through the ProteomeXchange Consortium via the PRIDE (https://www.ebi.ac.uk/pride/) partner repository with the data set identifiers for the DUBs profiling as PXD065005 ( **Username:** **Password:** NVTP9Qc8XBHX) and the Global protein ubiquitination data set as PXD065000 (**Username:** **Password:** uan2xgc8E6e6)

