## Supplementary document for "Bisphenol-A mediated ubiquitinome alteration triggers PPAR-alpha ubiquitination, affecting trophoblast cell migration"

### **Supplementary information:**

**Figure S1**: BPA-induced alteration in extravillous trophoblast cells.

**Figure S2**: Ubiquitinome profile of HTR8/SVneo cells.

**Figure S3**: In-silico study of BPA binding to PPAR-alpha.

**Figure S4**: Bisphenol-A influences PPAR-alpha ubiquitination and dysregulates its function.

**Table S1**: List of antibodies used in the experiments.

**Table S2**: List of gene specific primers used in the experiments.

**File S1**: List of activity-based protein profiling identified DUBs in HTR8/SVneo cells upon BPA treatment.

**File S2**: List of ubiquitinated proteins and peptides detected in control and BPA exposed condition.

**File S3**: MCODE clustered proteins from PPI enrichment and cluster identification of each MCODE component for BPA and Control condition.

**File S4**: List of enriched pathways from Metascape using unique ubiquitinated proteins from BPA and control groups.

**File S5**: list of curated substrate-DUBs-E3 Ligase connection based on Ub Browser 2.0 confidence score and Reactome pathways enriched of the common 33 substrate.

**File S6**: Overlap between PPAR-alpha, PPAR-gamma and PPAR-delta targets genes and KEGG pathway enrichment of unique PPAR-alpha targets (yellow highlighted).

**Supplementary Figures legends:**

**Figure S1: BPA induced alteration in extravillous trophoblast cells.** Quantification of wound area at different time points after 24-hour **A)** and 48-hour **B)** of incubation with 10 μM BPA. The area is plotted as mean ± SD (N=3). Statistical significance between different time-points was determined using two-way ANOVA, with *p <* 0*.*05 considered significant. Percentage of cells at healthy, early apoptotic, late apoptotic and necrotic states after treatment of BPA for 24 **C)** and 48 **D)** hours. Whole cell lysate of 24 and 48 hours of BPA treated and control sample subjected to western blot analysis for **E)** Total ubiquitin, **F)** Ubiquitin-Lys63, and **G)** Ubiquitin-Lys48. **H)** HA-tagged active ubiquitin probe (HA-UbVMe) was incubated with 24 or 48 hours of BPA treated and control Lysate followed by immunoprecipitation using anti-HA antibody to enrich active deubiquitinases visualized by the representative blot.

**Figure S2: Ubiquitinome profile of HTR8/SVneo cells A)** Unique Ub-modified protein enriched pathways from control group presented as networks of terms (each circle) connected with edges having similarity *>*0.3. **B)** Circle colour defines the associated cluster identity presented using bubble plot, where x axis and circle size shows protein count and log10(Qvalue) for each cluster Id, respectively. **C)** Protein-protein interaction network of unique ubiquitinated protein present in control group enriched from physical interactions in STRING (physical interaction score *>*0.132) and BioGrid. The MCODE algorithm enriched densely connected network components from the PPI network annotated using different colours. **D)** PPI network of identified proteins from MCODE defined coloured clusters and their associated functional description

**Figure S3: In-silico study of BPA binding to PPAR-alpha. A)** The table summarizes the modified peptides and ubiquitinated sites associated with 12 identified peptides of PPAR-alpha from the ubiquitinome profiling experiment. **B)** Predicted apo PPAR-alpha ligand-binding domain (LBD) structure (ice-blue) from Alphafold. **C)** BPA (green) docked into the ligand-binding pocket of PPAR-alpha Alphafold model. **D)** Crystal structure of PPAR-gamma LBD (tan) showing BPA (gray) bound in its experimentally determined binding site. **E)** Superposition of PPAR-alpha and PPAR-gamma highlighting the BPA binding site. The docked BPA pose in PPAR-alpha (green) overlaps the PPAR-gamma-bound BPA (gray) without steric clashes but displays a different orientation, suggesting subtle conformational differences in the binding pocket between the two receptors. **F)** Superposition of 69 PPAR-alpha ligand-binding domain (LBD) crystal structures reveals pronounced structural heterogeneity within the 242-267 loop region (dashed box), with positional displacements of up to ~15 Å among different conformations. This highlights the loop’s high flexibility and capacity to adopt multiple states. Zoomed view of the ligand-binding pocket showing the spatial arrangement of all key lysine residues K252, K257, K266, and K349 (colored sticks) involved in ubiquitination. **G)** Normal mode analysis (NMA) performed using the ProDy plugin in VMD shows ANM square fluctuations (Å²) for each residue. **H)** The 8HUK apo structure exhibits high fluctuations in the 242-267 loop, indicating intrinsic flexibility, whereas the BPA-bound Alphafold model displays reduced fluctuations in this region, reflecting ligand-induced stabilization. **I)** Ubiquitin (pink) docked into BPA bound PPAR-alpha shows bond length between K252, K257 and K266 and ubiquitin G76 suggesting a possible iso-peptide bond. **J)** The 8HUK apo structure docked with ubiquitin shows longer bond lengths between the PPAR-alpha Lysine and ubiquitin G76 indicating less possibility for an iso-peptide bond.

**Figure S4: BPA influences PPAR-alpha ubiquitination and dysregulates its function. A)** The integrated substrate-deubiquitinase-ligase network highlighting the PPAR-alpha node and its associated DUBs and E3 ligases. The edge width denotes the confidence score. **B)** Venn diagram showing overlap between the PPAR-alpha (blue), PPAR-gamma (green) and PPAR-delta (red) target genes curated from TFLink database. Red square highlights the 41 unique PPAR-alpha targets used for STRING PPI enrichment analysis. **C)** KEGG pathway enrichment of 41 unique PPAR-alpha targets shows top 10 pathways where circle size and colour indicate the gene count and -log10(FDR) respectively. The fold enrichment of respective pathway is plotted in x-axis. **D)** The quantified intensity of ubiquitin PPAR-alpha colocalised puncta in the nucleus of HTR8/SVneo cells after BPA exposure. The measured gene expression of **E)** IL6R and **F)** IGFBP1 relative to GAPDH in BPA treated and control samples. Mean ± SEM (N=3) values are reported based on three independent biological replicates. Statistical comparisons were conducted using student’s t-test, with significance defined as *p* < 0.05.

**Table S1**:

| **Species** | **Antigen** | **Cat#, Company** | **Dilution** | **Application** |
| --- | --- | --- | --- | --- |
| Rabbit | MMP-2 | 40994, Cell Signaling | 1:1000 | WB |
| Rabbit | Beta-Actin | 4970, Cell Signaling | 1:5000 | WB |
| Rabbit | MMP-9 | ab38898, Abcam | 1:1000 | WB |
| Mouse | Ubiquitin | sc-271289, Santa Cruz | 1:100 | WB, IF |
| Rabbit | Ubiquitin-Lys63 | 05-1308, Merck | 1:1000 | WB |
| Rabbit | Ubiquitin-Lys48 | 05-1307, Merck | 1:1000 | WB |
| Rabbit | HA | RM305, Invitrogen | 1:1000 | WB |
| Mouse | GAPDH | 39-8600, Invitrogen | 1:5000 | WB |
| Rabbit | PPAR-alpha | PA1-822A, Invitrogen | 1:1000, 1:100 | WB, IF, IP |
| Rabbit | MuRF1 | PA5-76695, Invitrogen | 1:2000 | WB |
| Rabbit | TRIM46 | 21026-1-AP, Proteintech | 1:1000 | WB |
| Rabbit | USP5 | 10473-1-AP, Proteintech | 1:1000 | WB |
| Rabbit | UCHL1 | 14730-1-AP, Proteintech | 1:2000 | WB |

**Table S2**:

| **Target Gene** | **Forward (5’ to 3’)** | **Reverse (5’ to 3’)** |
| --- | --- | --- |
| MuRF1 | CTTCCAGGCTGCAAATCCCTA | ACACTCCGTGACGATCCATGA |
| TRIM46 | TTCCGACCCAAGGGCCTTAT | AGAGTTGACATACCAGGCGTT |
| HUWE1 | TTGGACCGCTTCGATGGAATA | TGAAGTTCAACACAGCCAAGAG |
| UCHL1 | CCGAGATGCTGAACAAAG | CAGAGACTCCTCTTCCAG |
| USP5 | CGGCCAGCGAGTCTACTTG | AAGGTCAAATCCGCCTTCAAC |
| TGFB1 | GAGCCTGAGGCCGACTACTAC | CACGTGCTGCTCCACTTTTA |
| IGFBP1 | TTTTACCTGCCAAACTGCAACA | CCCATTCCAAGGGTAGACGC |
| CPT1A | TCCAGTTGGCTTATCGTGGTG | TCCAGAGTCCGATTGATTTTTGC |
| CITED2 | AACCAGCACTTCCGAGATTGC | AATCAGTGTCTATGACATTGGGC |
| CYP1A1 | TGGCATCCTCTACAGACTCCTG | CTTCAGGTTGCGTGCCATCTCA |
| IL6R | CCCCTCAGCAATGTTGTTTGT | CTCCGGGACTGCTAACTGG |
| STAT5A | GCAGAGTCCGTGACAGAGG | CCACAGGTAGGGACAGAGTCT |
| HMGCS1 | CATTAGACCGCTGCTATTCTGTC | TTCAGCAACATCCGAGCTAGA |
| CGB5/CGB3 | TGAGCCACTCCTGCGCCC | CAGCCCCTGGAACATCTCCA |
| NFKBIA | CTCCGAGACTTTCGAGGAAATAC | GCCATTGTAGTTGGTAGCCTTCA |
| GAPDH | GCACCGTCAAGGCTGAGAAC | TGGTGAAGACGCCAGTGGA |

**Figure S1:**

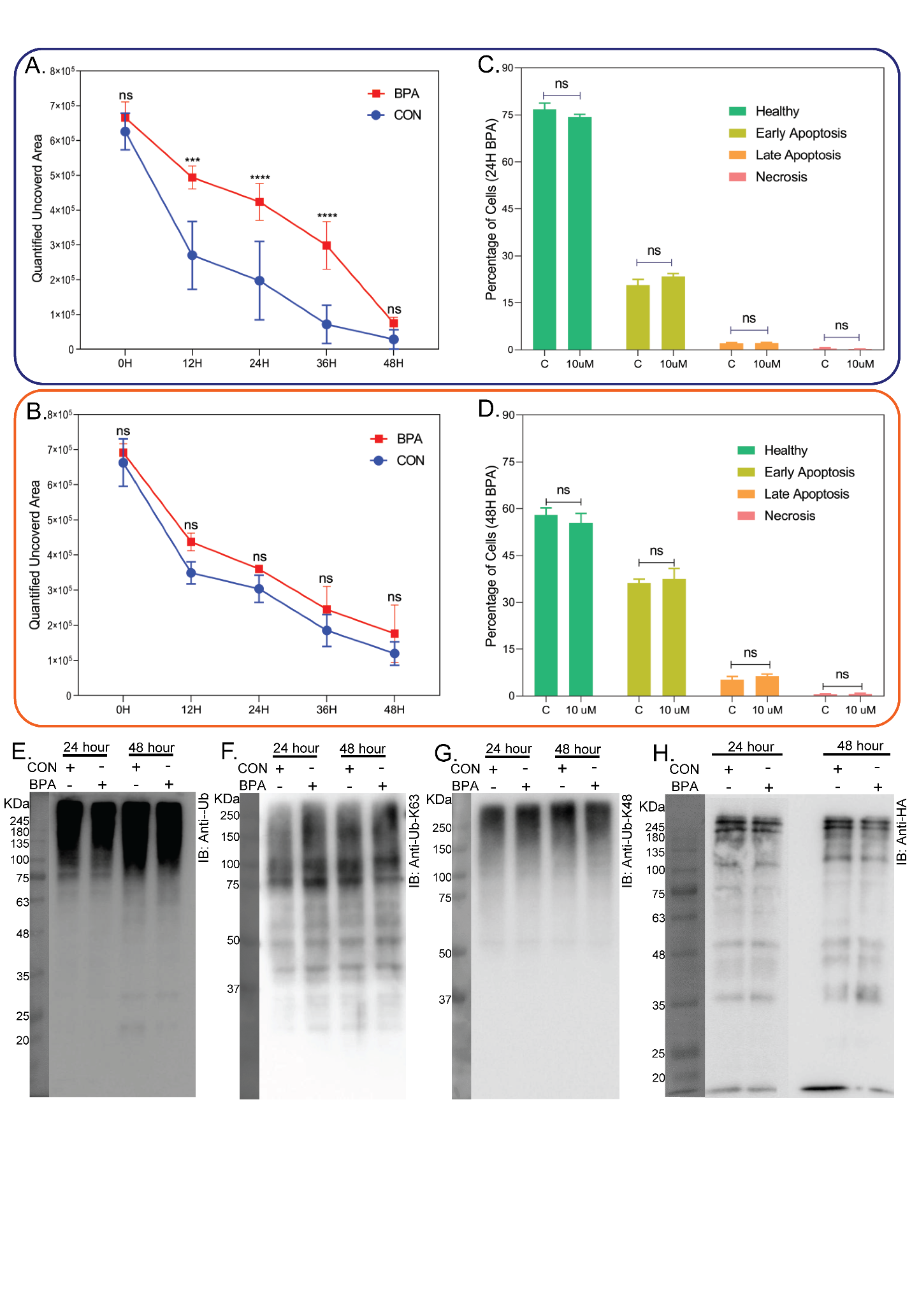

**Figure S2:**

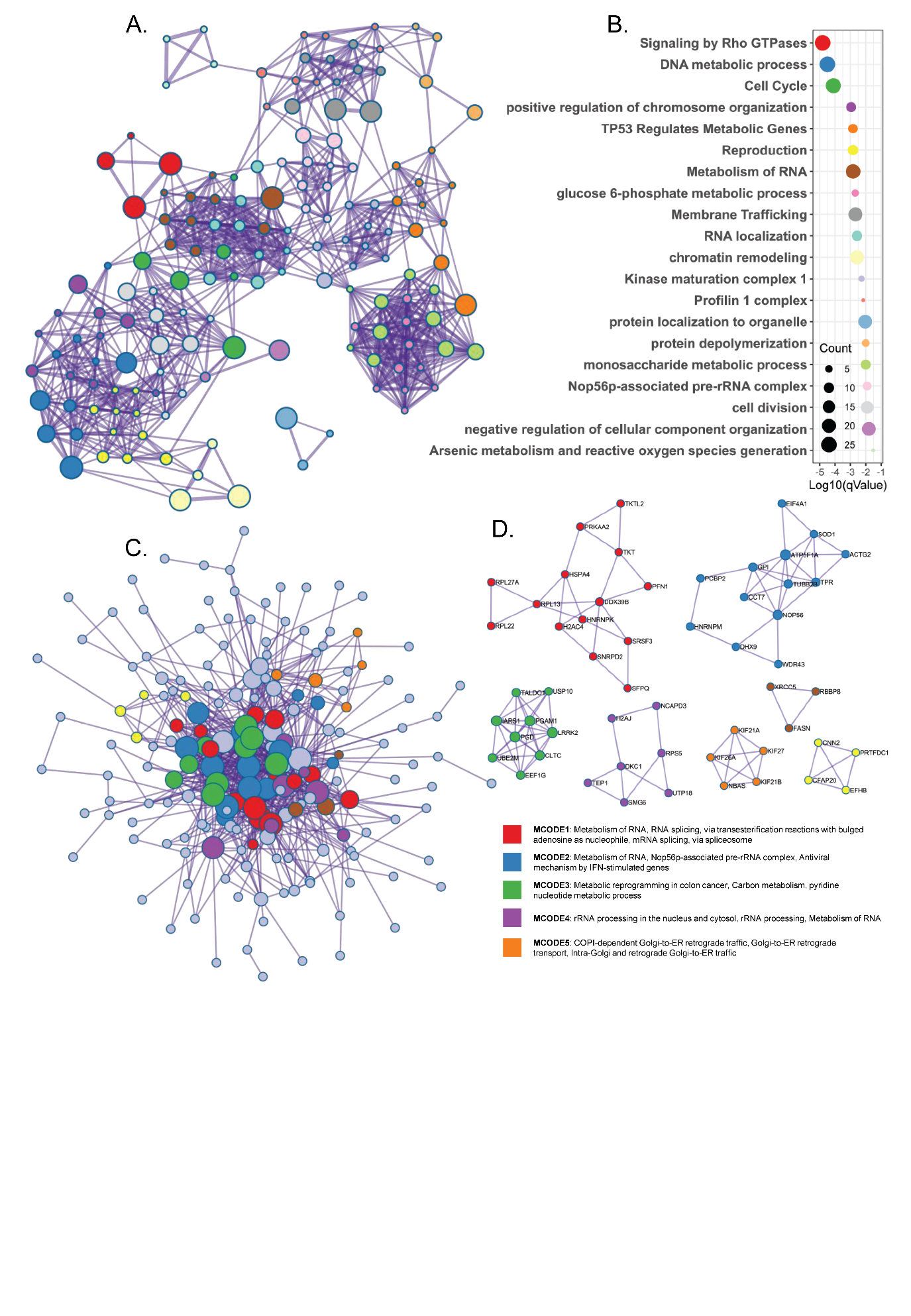

**Figure S3:**

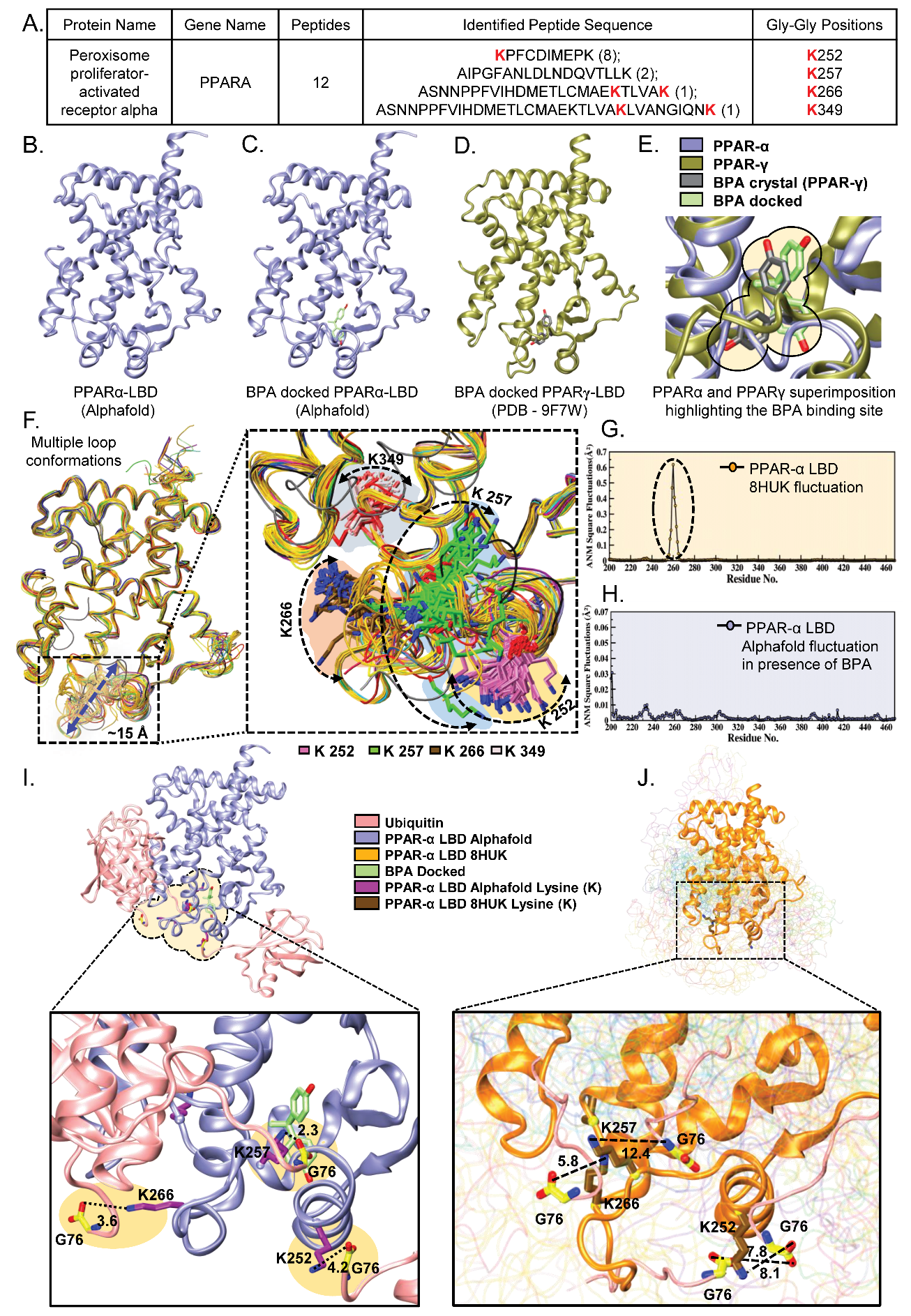

**Figure S4:**

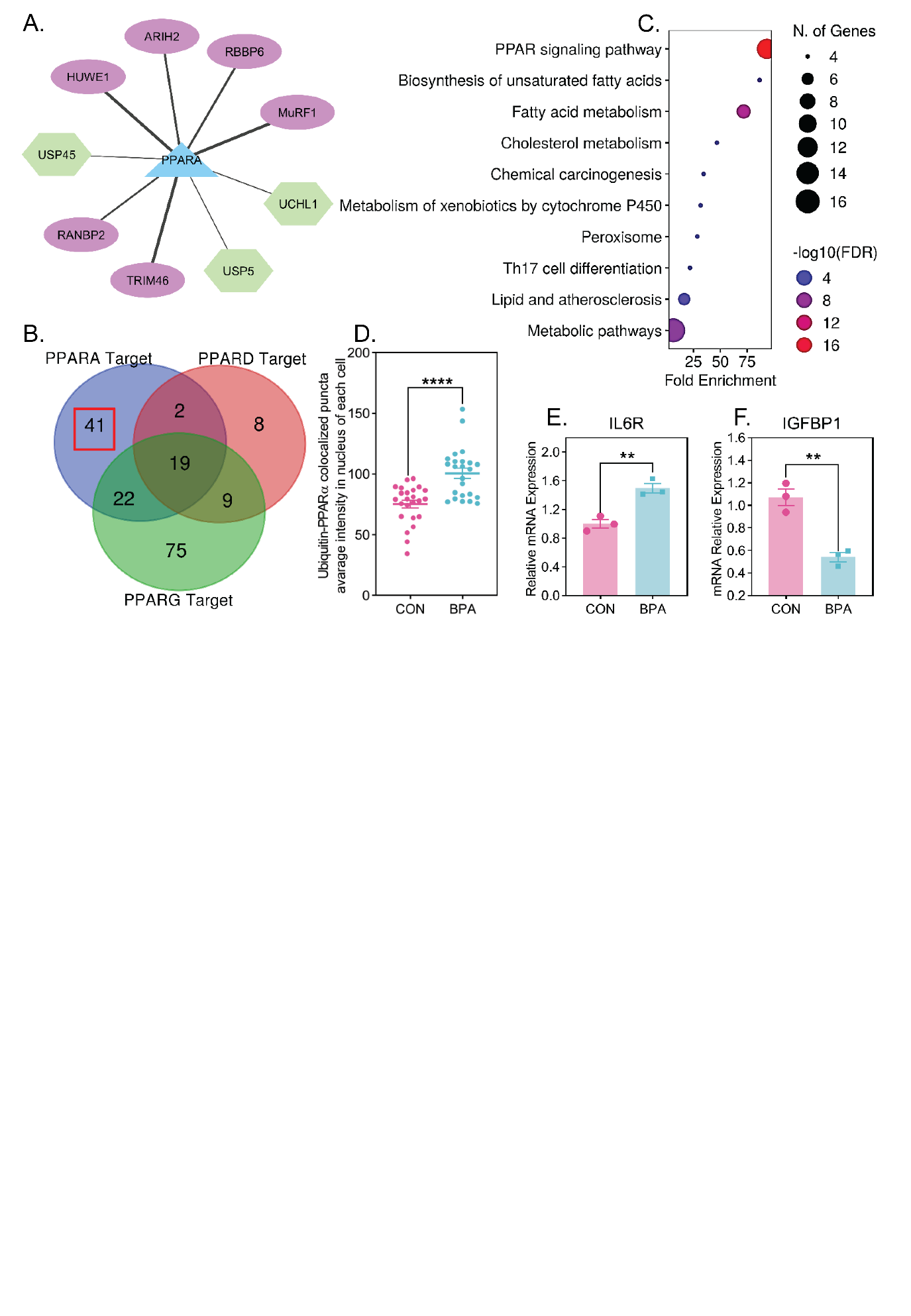
